# Addressing pandemic-wide systematic errors in the SARS-CoV-2 phylogeny

**DOI:** 10.1101/2024.04.29.591666

**Authors:** Martin Hunt, Angie S. Hinrichs, Daniel Anderson, Lily Karim, Bethany L Dearlove, Jeff Knaggs, Bede Constantinides, Philip W. Fowler, Gillian Rodger, Teresa Street, Sheila Lumley, Hermione Webster, Theo Sanderson, Christopher Ruis, Benjamin Kotzen, Nicola de Maio, Lucas N. Amenga-Etego, Dominic S. Y. Amuzu, Martin Avaro, Gordon A. Awandare, Reuben Ayivor-Djanie, Timothy Barkham, Matthew Bashton, Elizabeth M Batty, Yaw Bediako, Denise De Belder, Estefania Benedetti, Andreas Bergthaler, Stefan A. Boers, Josefina Campos, Rosina Afua Ampomah Carr, Yuan Yi Constance Chen, Facundo Cuba, Maria Elena Dattero, Wanwisa Dejnirattisai, Alexander Dilthey, Kwabena Obeng Duedu, Lukas Endler, Ilka Engelmann, Ngiambudulu M. Francisco, Jonas Fuchs, Etienne Z. Gnimpieba, Soraya Groc, Jones Gyamfi, Dennis Heemskerk, Torsten Houwaart, Nei-yuan Hsiao, Matthew Huska, Martin Hölzer, Arash Iranzadeh, Hanna Jarva, Chandima Jeewandara, Bani Jolly, Rageema Joseph, Ravi Kant, Karrie Ko Kwan Ki, Satu Kurkela, Maija Lappalainen, Marie Lataretu, Jacob Lemieux, Chang Liu, Gathsaurie Neelika Malavige, Tapfumanei Mashe, Juthathip Mongkolsapaya, Brigitte Montes, Jose Arturo Molina Mora, Collins M. Morang’a, Bernard Mvula, Niranjan Nagarajan, Andrew Nelson, Joyce M. Ngoi, Joana Paula da Paixão, Marcus Panning, Tomas Poklepovich, Peter K. Quashie, Diyanath Ranasinghe, Mara Russo, James Emmanuel San, Nicholas D. Sanderson, Vinod Scaria, Gavin Screaton, October Michael Sessions, Tarja Sironen, Abay Sisay, Darren Smith, Teemu Smura, Piyada Supasa, Chayaporn Suphavilai, Jeremy Swann, Houriiyah Tegally, Bryan Tegomoh, Olli Vapalahti, Andreas Walker, Robert J Wilkinson, Carolyn Williamson, Xavier Zair, IMSSC2 Laboratory Network Consortium, Tulio de Oliveira, Timothy EA Peto, Derrick Crook, Russell Corbett-Detig, Zamin Iqbal

## Abstract

The SARS-CoV-2 genome occupies a unique place in infection biology – it is the most highly sequenced genome on earth (making up over 20% of public sequencing datasets) with fine scale information on sampling date and geography, and has been subject to unprecedented intense analysis. As a result, these phylogenetic data are an incredibly valuable resource for science and public health. However, the vast majority of the data was sequenced by tiling amplicons across the full genome, with amplicon schemes that changed over the pandemic as mutations in the viral genome interacted with primer binding sites. In combination with the disparate set of genome assembly workflows and lack of consistent quality control (QC) processes, the current genomes have many systematic errors that have evolved with the virus and amplicon schemes. These errors have significant impacts on the phylogeny, and therefore over the last few years, many thousands of hours of researchers time has been spent in “eyeballing” trees, looking for artefacts, and then patching the tree.

Given the huge value of this dataset, we therefore set out to reprocess the complete set of public raw sequence data in a rigorous amplicon-aware manner, and build a cleaner phylogeny. Here we provide a global tree of 4,471,579 samples, built from a consistently assembled set of high quality consensus sequences from all available public data as of June 2024, viewable at https://viridian.taxonium.org. Each genome was constructed using a novel assembly tool called Viridian (https://github.com/iqbal-lab-org/viridian), developed specifically to process amplicon sequence data, eliminating artefactual errors and mask the genome at low quality positions. We provide simulation and empirical validation of the methodology, and quantify the improvement in the phylogeny. We hope the tree, consensus sequences and Viridian will be a valuable resource for researchers.

## Introduction

On the eve of the SARS-CoV-2 pandemic, had one commissioned a poll of phylogeneticists on whether their methods were adequate for current public health needs, the overall response would have been in the affirmative. At that point, most people were analysing relatively small datasets (*N<*5000), usually carefully curated and generally studied by people working closely with those obtaining and processing the clinical samples, or indirectly, via national public health organisations. Data were usually small, clean, and there was limited urgency. One year later, all of these statements would no longer be true. The SARS-CoV-2 pandemic placed unprecedented strains on the genomics and bioinformatics communities in terms of scale, turnaround time, and coordination. In every dimension, tools and systems were pushed far beyond expectations. Despite significant efforts and innovations, numerous steps in the process (i.e. from patient to global phylogenies and dashboards) required prioritizing speed and practicality over absolute accuracy. This was the right thing to do at the time as it enabled real-time management decisions to be taken. However, since there was no unified genome assembly or QC process, the end result has been that the set of SARS-CoV-2 genomes, on which future evolutionary and vaccine analyses will be based, contain a large number of systematic errors [1, 2]. The goal of this study is to re-assemble all publicly available SARS-CoV-2 raw sequence data with a single analysis workflow to remove the vast majority of these errors, thereby building a higher quality phylogenetic tree for all our benefit.

Unlike the sequencing of bacterial genomes after culture (where the details of sequencing and assembly can stay the same over reasonably long periods) the specifics of viral sequencing and assembly during the pandemic had to keep changing, as we describe below. This resulted in a myriad of inconsistencies across the globe, and errors in consensus sequences. A fundamental constraint on sequencing of SARS-CoV-2 was the fact that viral load in patient samples was generally very low and highly variable, as a result of which the most common way to sequence was via tiled amplicons (as had been done previously for other viruses [3]). Here, the genome is divided into overlapping “tiles”, each of which is independently PCR-amplified, guided by PCR primers at either end of the tile. That this was possible at all was thanks to two things: the early release of the genome sequence [4, 5], and Quick et al’s rapid production of a set of primers, the first “ARTIC” (acronym referring to a consortium) primer scheme [6]. A feature of any tiled amplicon scheme is that, as the virus evolves, eventually mutations within primer-binding sites will lead to failed amplification of the associated tile, creating gaps in the genome sequence data (“dropouts”). This is to be expected and necessitates the development of an updated scheme with new primers. Additionally, many genome assembly software pipelines implicitly made the false assumption that in the absence of data (no reads from an amplicon) one should infer the sequence as being that of the reference genome, which in the case of SARS-CoV-2 is also the ancestral sequence. Thus at various points during the pandemic, researchers analysing the phylogeny would find a sudden crop of genomes “reverting to the ancestor”.

In Figure 1a we show part of a tree with the leaves coloured to show what base that genome has at a specific position – purple for the ancestral base, and green for the derived (new) base caused by a mutation shown as a white star. One single mutation explains that data. In Figure 1b, we show the impact of wrongly assigning the ancestral base at the lowest-but-one leaf (fourth purple down). Here, the most parsimonious way to explain this is with a second mutation (red star) “reverting” back to the ancestral purple. In Figure 1c we show part of the global SARS-CoV-2 phylogeny hosted at taxonium.org (accessed 9^th^ April 2024), zoomed in to show where Omicron branches from the ancestor. Leaves are coloured by the genotype of genome position 22813 (codon 417) in the spike gene (again purple is ancestral). In the blow-up we see within the green (Omicron) clade, a striking spray of purple that does not sit cleanly in any subclade. Patterns like this are in general more likely to be due to assembly artefact than multiple independent reversions. Such errors can have considerable impact on our inferences about the underlying biology – in this case K417N is a mutation that affects antibody escape [7], and systematic errors like this can lead to misinterpretation. However, although one can use a reversion count as a metric of whether we suspect there are assembly problems, reversions are not always errors. For example SARS-CoV-2 has a C to T mutation bias [8, 9] (strictly a C to U, as it is an RNA virus, but we convert to DNA space for phylogenetics), so if you have a T to C mutation on a phylogenetic branch leading to a large clade, you may expect to see multiple reversions back to T in that clade.

**Figure 1:**
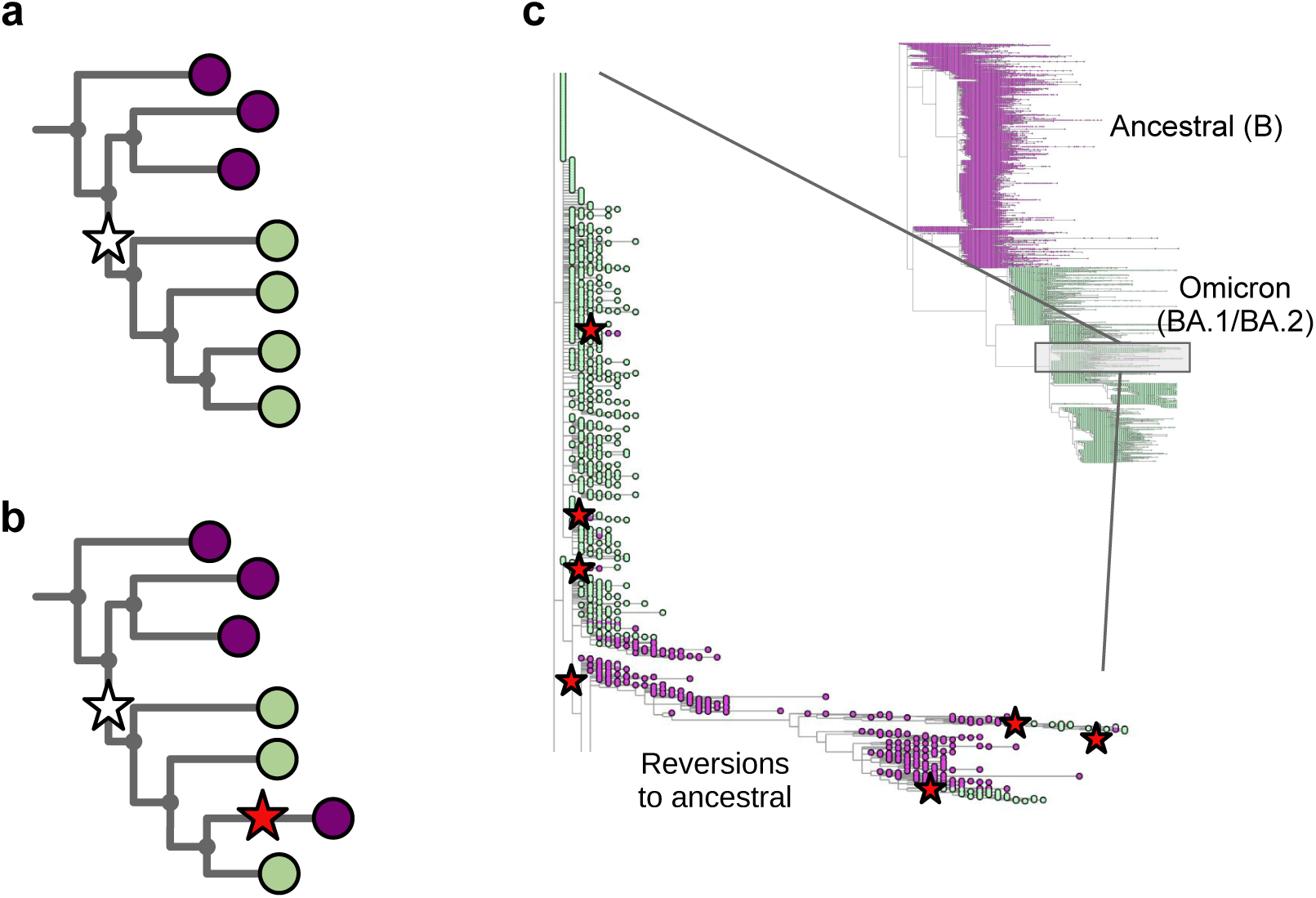
Assemblers which wrongly default to the reference base in the absence of data cause reversions in the phylogeny. **a)** Cartoon phylogeny built from perfect genomes, with leaves coloured by genotype at a specific position X (purple – ancestral base, green – derived base). Just one mutation at this site, shown as a white star, is needed to explain the data. **b)** Cartoon showing the effect of assembly software assuming that a genome is identical to the reference genome when there is no data – here the amplicon containing position X is dropped in the lowest-but-one genome on the tree, creating one lone purple leaf. The tool which infers the phylogeny looks for a parsimonious explanation for this colour distribution, and concludes it was caused by a mutation (white star) followed by a “reversion” back to the ancestral base (red star). Errors in assembly caused by reference-bias tend to create enrichments of reversions. **c)** Part of the current UShER SARS-CoV-2 phylogeny, coloured by genotype at genome position 22813 (spike codon 417). Blow-up shows multiple reversions back to the ancestral purple. A non-exhaustive set of artefactual mutations (reversions, unreversions, re-reversions etc) are shown with red stars, where there is a flip back and forth from green to/from purple.

There are a number of other possible technical artefacts that can arise (e.g. primer dimers [10], interactions between amplicons [17], or primers binding in non-canonical sites [15]) which should be expected and handled, otherwise additional errors will result. Unfortunately, these errors often correlated with individual sequencing centres, which themselves correlated with local prevalence of particular lineages at particular times. In addition, where amplicon dropout was incomplete, the likelihood of wrongly imputing the reference genome at a particular position becomes a function of decreasing amounts of sample RNA, creating a false relationship between genotype and viral load [14].

Because of amplicon dropouts, as the pandemic progressed and sequential waves of Variants of Concern (VOCs) arose, the ARTIC primer scheme was updated multiple times to restore amplification, as well as a slew of alternative options (Midnight [18], AmpliSeq (Thermo Fisher Scientific), VarSkip (https://github.com/nebiolabs/VarSkip, etc). Each VOC wave brought mutations in primer bindings sites leading to amplicon dropouts, and a subsequent wave of artefacts in genomes as these were mishandled (see Figure 2). New amplicon schemes were then introduced, and gradually taken up, solving previous dropout problems, but also followed by smaller waves of new artefacts in the genomes, sometimes caused by primers not being correctly trimmed and being incorporated into assemblies. It is no exaggeration to say that since this issue was first raised [2], thousands of person-hours of time have been spent manually looking through trees and genomes trying to decide if strange phenomena are artefacts or not. Some of us (RCD, AH) have been maintaining the global phylogenetic tree of SARS-CoV-2 since 2021 [19], and the only way we have been able to maintain the integrity of the tree has been to a) completely mask 150 nucleotide positions in the genome, as they are systematically too often wrong to ever be trusted, and b) systematically mask (ie ignore) certain mutations on specific branches of the tree. As artefacts ebbed and flowed, and were discovered by analysts, the masking had to be updated (see Figure 2 and Supplementary Figure S1). After the mammoth global efforts to sequence and collate these SARS-CoV-2 genomes, the richest dataset of any pathogen to date, it is critical to now reprocess and clean this data, providing a firm foundation for future discoveries.

**Figure 2:**
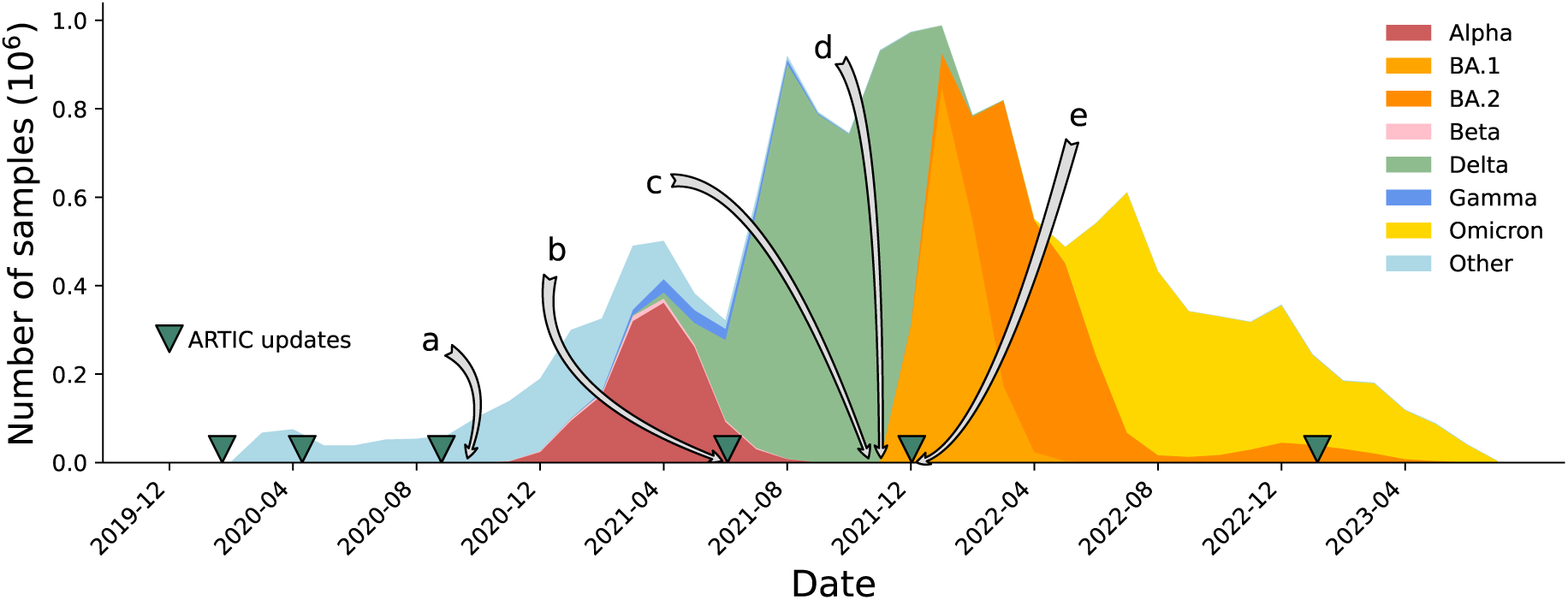
Timeline of the SARS-CoV-2 pandemic from December 2019 to July 2023, with selected events relating to problems with sequencing and consensus calling labelled a-e. Releases of ARTIC primers schemes (versions 1, 2, 3, 4, 4.1, 5.3.2) are marked with green triangles. a) Primer dimers cause amplicon dropouts [10] and 28% of GISAID [11] sequences deposited in September 2020 have at least one gap of length at least 200bp [12]. b) A 9bp deletion in the primer binding region of ARTIC V3 amplicon 73 causes missing data [13]. c) Dropouts causing artefacts at Spike 95 and 142 [14]. d) ARTIC v4 roll out triggers artifactual mutations in some pipelines [15]. e) Omicron samples cause ARTIC v4 amplicon dropout, triggering the update to ARTIC v4.1 [16].

As of June 2024, there were approximately 6 million SARS-CoV-2 raw sequence datasets deposited in the SRA/ENA, very few of which had metadata recording the primer scheme and the assembly pipeline used (data from COG-UK being a notable but geographically localised exception). In this paper we will describe our amplicon-aware assembly and QC processes, reprocessed these genomes and measured the improvements in the genomes and phylogeny, and provide these data as a resource for the whole community.

## Results

We set out to reprocess all available SARS-CoV-2 sequence read data, generating new consensus genomes through an assembly workflow designed for tiled amplicon schemes with a rigorous quality-control process, and thereby build a global phylogeny that minimizes the need for masking unreliable parts of the genome and tree.

To this end, we created Viridian, an efficient amplicon-aware assembler to consistently handle Illumina, Oxford Nanopore, and Ion Torrent reads. Since publicly shared sequence data does not generally have metadata logging the primer scheme used, Viridian first identifies the amplicon scheme from the input reads. In light of this, with knowledge of where primers bind, it then makes consensus sequences for each amplicon by building a partial-order alignment graph of the reads using Racon [20], an approach which will detect indels more robustly than one based on pileups. Viridian then merges the per-amplicon consensuses into a single consensus and calls variants. To evaluate the confidence of each position in this consensus, it remaps the reads to the consensus, identifies unsupported positions, and using this, finally outputs a high quality sequence that has low quality bases masked. The emphasis throughout is on minimising errors, in particular where amplicon primers bind, producing a consensus sequence where all unmasked positions should be correct.

We performed three evaluations of Viridian against two existing ARTIC workflow implementations: ARTIC-ILM (for Illumina) and ARTIC-ONT (for Nanopore) (see Methods). The data used were 1) simulated data, 2) a “truth set” of 67 runs from 27 isolates with known results, and 3) a larger dataset (*N* =12287, “Early Omicron”) from multiple countries in Africa from November 2021 to March 2022 that includes the emergence of the Omicron variant.

### Primer Scheme identification

We first evaluated our method for identifying primer schemes (see Methods) using two datasets where we knew the correct primer scheme – these consisted of 8,000 simulated genomes and 67 curated truth genomes. There were zero errors. We then used 2,341,118 Illumina and 122,410 Oxford Nanopore samples where the ENA/SRA metadata had an ARTIC primer scheme version entry of 3 or 4, and compared with the call from Viridian (Supplementary tables 1,2). There was 99.7% agreement for Illumina and 98.2% for Oxford Nanopore samples. A manual investigation of a subset (*N* =20) of the discordances concluded that the remaining errors were likely metadata errors in the ENA/SRA: in 19/20 cases, the pileups were categorical that Viridian was correct, and in the remaining one, the data were inconclusive (supplementary text and Figures S2-6). Note that both the truth set and the ENA/SRA data contain samples where tagmentation during the library preparation caused fragmented reads, confirming the method worked there too.

### Simulations

We simulated a SARS-CoV-2 tree of 8,000 genomes, including SNP errors in primers and amplicon dropouts. Illumina and Nanopore reads were simulated from each genome, from simulated amplicons using the ARTIC v4 scheme. To evaluate the accuracy of resulting consensus sequences from ARTIC-ILM, ARTIC-ONT, and Viridian, a novel pipeline was developed called CTE (covid truth evaluation, see Methods), which evaluates each consensus sequence using the truth to classify each position in the genome as correct or as an error. Results were highly consistent across all tools and amplicon schemes (Supplementary Tables S3a-d). Although there were overall very few errors, ARTIC-ONT had notably more indel errors than Viridian (Supplementary Tables S3c,d).

### Empirical truth dataset

The tools were compared on a truth dataset of 67 high quality sequencing runs from 28 samples, comprising a mix of Illumina and Nanopore reads, and ARTIC (v3, v4, v4.1) and Midnight amplicon schemes. The truth, including all expected SNPs in all runs, was determined by manual inspection of reads mapped to the reference genome. Similarly to the simulations, all tools performed well, with few errors (Supplementary Tables S4,5), and Viridian performing better with respect to indels on Nanopore data (Supplementary Table S5e,f). We measured the peak RAM and total CPU time of each truth set run. Viridian had mean peak RAM usage of 444MB and mean CPU time 154s, whereas ARTIC-ILM and ARTIC-ONT used 1.45GB of RAM and took 366s, and 1.80GB of RAM, and took 561s respectively (Supplementary Table S6, Supplementary Figure S7).

### African “Early Omicron” dataset

Next we evaluated our own empirical dataset, sequenced and assembled at CERI in South Africa, with samples from November 2021 to March 2022, including Variants of Concern Alpha, Beta and Delta, and also encompassing the emergence of the Omicron variant. The 12,287 samples were from South Africa (*N* =8,645), Angola (*N* =957), Mozambique (*N* =619), Mauritius (*N* =488), Malawi (*N* =480), Cameroon (*N* =344), Zimbabwe (*N* =333), Ethiopia (*N* =232), Uganda (*N* =102), Namibia (*N* =83) (and 4 with unknown country), and include Illumina (*N* =9,935) and Nanopore (*N* =2,352) runs, using either ARTIC (*N* =11,070 including versions 3,4, and 4.1) or Midnight (*N* =1,217) amplicon schemes (Supplementary Table S7). Each sample was processed with Viridian and ARTIC-ILM/ARTIC-ONT as appropriate, and the results compared with our original assemblies [21] which have previously been shared to the UShER [22, 23] SARS-CoV-2 phylogeny via GISAID. We scanned all positions in all consensus assemblies for “hard errors”, where the majority of the reads disagreed with the consensus (for example the consensus called an A but most reads say G, see Methods). We found systematic positional errors (which were specific to primer scheme and sequencing technology) in the original consensuses and the ARTIC-ONT assemblies. The errors were significantly reduced in the ARTIC-ILM workflow although some did remain. By contrast the errors were completely removed by Viridian. This is summarised in Figure 3.

**Figure 3:**
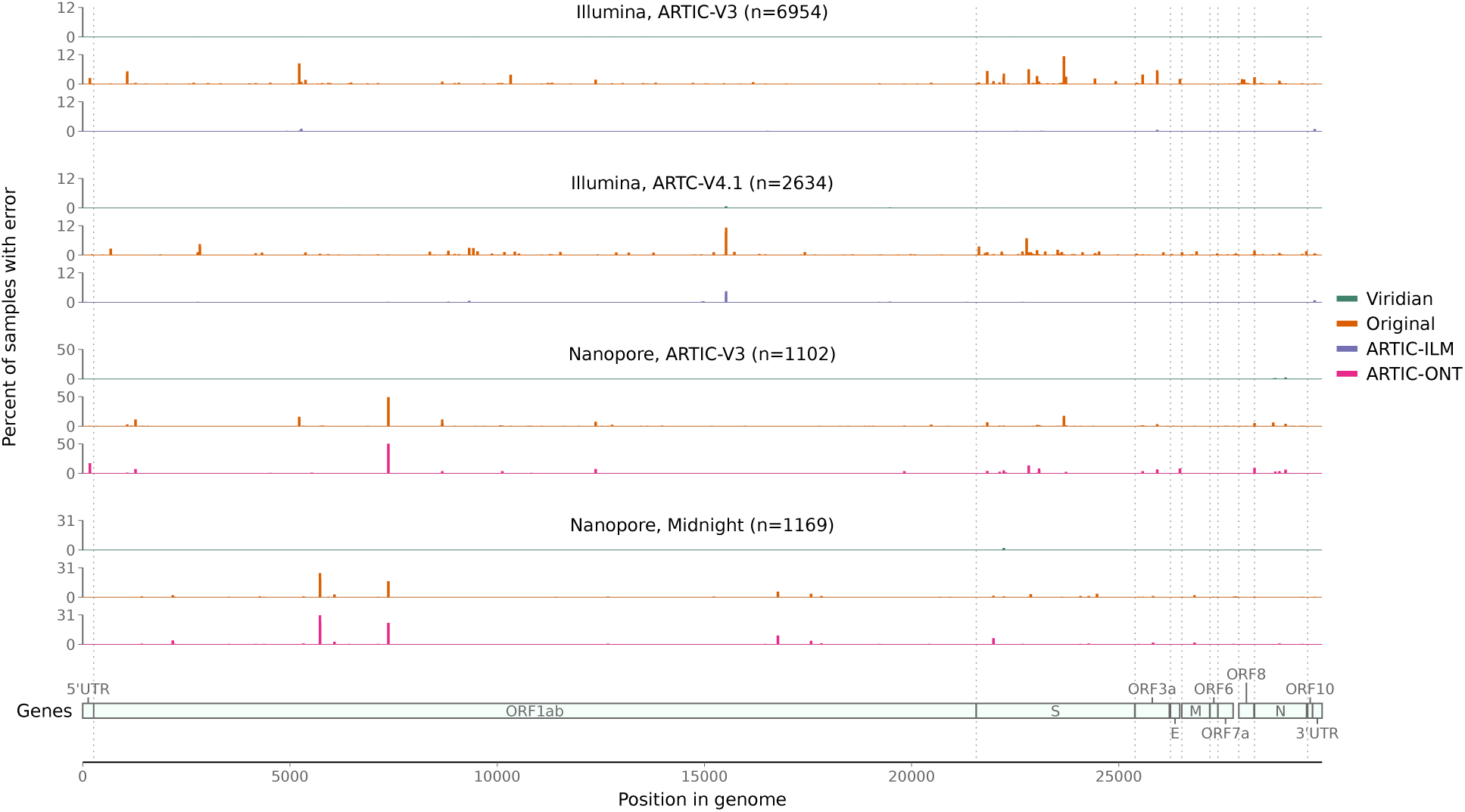
Errors across the genome in consensus sequences from the “Early Omicron” African dataset, split by sequencing technology and amplicon scheme. Plots show the percent of consensus sequences with an error (*y*-axis), taking the maximum value in windows of length 50bp (*x*-axis). Error here is defined as where the consensus sequence has an A/C/G/T call, the read depth passes Viridian’s default filters (see methods), and the reads support a different A/C/G/T call. Results are shown for Viridian, the original assemblies, and for the ARTIC-ILM and ARTIC-ONT assembly workflows.

### Assembly and evaluation of the global data

We processed all Illumina, Nanopore and Ion Torrent SARS-CoV-2 sequencing runs from the ENA/SRA as of 2^nd^ March 2023, keeping all 3,960,704 that passed QC (see Methods) and produced a consensus sequence using Viridian. We also obtained all matching entries from GenBank, giving an “intersection set” of 3,311,456 samples with both a Viridian and GenBank consensus sequence. We then built a tree of each of these three data sets – all 3,960,704 Viridian sequences, Intersection/Viridian (i.e. the Viridian assemblies of the intersection set), and Intersection/GenBank (i.e. the GenBank assemblies of the intersection set) – using MAFFT [24] and UShER (reverting deletions to the ancestral sequence and excluding insertions, see Methods). Note that these trees

a) are built from unmasked consensus genomes, unlike the current UShER global SARS-CoV-2 phylogeny, which pre-masks a list of “Problematic Sites” in the genome where the community has determined assemblies may be unreliable, and
b) do not have any forcible masking of particular mutations on the branches of specific Variants of Concern, unlike the current public SARS-CoV-2 tree. To assess the improvement in accuracy of a tree built from Viridian sequences, we next compared the Viridian and GenBank intersection set trees.

### Ns and Pango assignment

A scatterplot comparing the number of Ns in the Viridian versus GenBank assemblies (Supplementary HTML file) showed very little correlation, and a strong enrichment of points where there were many more Ns in the Viridian assembly – *N* =1,604,389 (53.4%) of GenBank assemblies had no Ns, compared to *N* =1,197,638 (39.8%) of Viridian assemblies. There were more Ns in the GenBank assembly for 9% of samples versus 49% samples with more Ns in the Viridian assembly; of those samples with more Ns in the Viridian assembly, 29% had zero Ns in the GenBank assembly. This is consistent with the known issue that for some software pipelines, portions of the reference sequence had been used to fill in dropouts for a large number of sequences, and this effect alone will have been a significant cause of reversions in the tree. Nevertheless, analysis at the lineage level using Pangolin showed very strong agreement, with only 0.98% (*N* =29,475) of samples having discordant assignments. Of the mismatches, the majority (77%) were parentchild, with Viridian assembly the child (i.e. more specific) in 60% of those. Only 0.01% (N=287) mismatched at the variant level. No Viridian assembly was “Unassigned”, compared with 87 of the GenBank assemblies. Analysis of the results by collection date, country, technology and primer scheme revealed no category enriched for disagreements.

### Indel calls

In samples where Viridian and GenBank assemblies result in the same Pangolin variant, indel calls are generally concordant and either very dominant or very rare. The characterising insertion of TAC after position 21990 (S:YY144–145TSN) in Mu is an exception, found in 90% of Viridian assemblies but only 60% of GenBank assemblies. In samples where Viridian/GenBank have mismatched WHO variant calls, we see fewer indels per sample in GenBank versus Viridian (Supplementary HTML File). Notable differences at variant-defining indel sites – in particular, for samples assigned Delta for the Viridian assembly and Omicron for the GenBank assembly, we see two Delta-defining indels that are present in the Viridian assemblies, but absent in the GenBank assemblies. We show in Supplementary Figure S8 those positions where there is discordance between Viridian and GenBank.

### Reversions

One of the key signals of artefactual problems used during the pandemic, was finding positions in the genome (or branches of the tree) with very large numbers of reversions. We therefore used Matutils [19] and custom scripts to count the number of reversions in both trees, and plot this in two ways. In Figure 4a, we show one minus the cumulative density function of reversions in the two trees, showing that the Viridian tree has far fewer positions with many reversions. To understand which positions are problematic, in Figure 4b we show a scatter plot comparing number of reversions at each position of the genome, in the Viridian and GenBank trees, with a blow-up of the central region in Figure 4c. The main issue for phylogenetic analysis is positions with large numbers of reversions, so we care more about the graph away from the origin. We see that apart from a handful of positions far to the right and below the line *y*=*x*, all positions have fewer reversions in the Viridian tree. In other words, a smaller set of positions can be masked in the Viridian tree than in the GenBank tree in order to greatly reduce the number of reversions. For example, the Genbank tree has 63 positions with 200 or more reversions, while the Viridian tree has only 20. See also Supplementary Figure S9 for the specific example of genome position 22813 (introduced earlier in Figure 1), comparing the current UShER global phylogeny with the Viridian tree.

**Figure 4:**
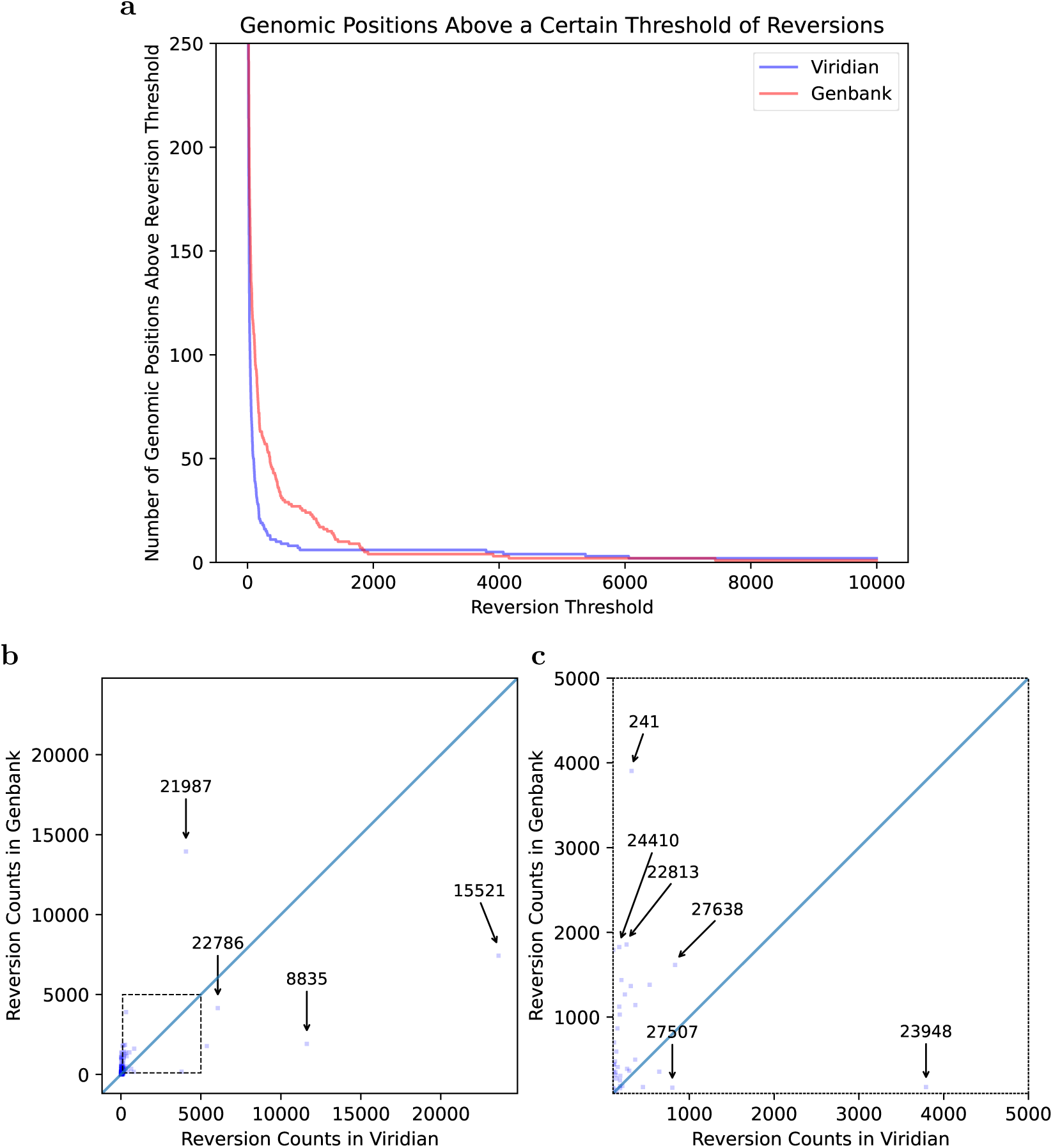
Most variable sites cause fewer reversions in the Viridian tree than the GenBank tree. a) Plot showing how many positions in the genome (y axis) have at least *N* reversions (x axis) in each tree (Viridian in blue, GenBank in red). Viridian curve drops faster, having fewer positions that create many reversions. b) Scatterplot comparing count of reversion mutations found in GenBank Dataset (*y* axis) and Viridian dataset (*x* axis). Note (0,0) is slightly indented from the origin of the plot. Each point represents a position of the SARS-CoV-2 genome. Three points below the line *y*=*x* are highlighted (labelled by genomic coordinates: 22786, 8835, 15521) where Viridian has particularly high numbers of reversions, and one (labelled 21987) for GenBank. c) Blow up of dotted square from panel b) showing vast majority of variable sites in the genome lie above the line y=x.

### Improved accuracy of lineage growth rate estimates

We ran Py*R*_0_, a hierarchical bayesian regression model that measures growth rates of SARS-CoV-2 lineages using genetic, temporal and geographical data [25]. When we ran this model on the Viridian tree, precision improved more than threefold on average compared with running the model on a Genbank tree. B– and BA-descended lineages had the largest decrease in the uncertainty of their growth rate measurements (see Figure 5). Improvements in precision occurred while maintaining accuracy. Supplementary Figures S10-12 provide more detail.

**Figure 5:**
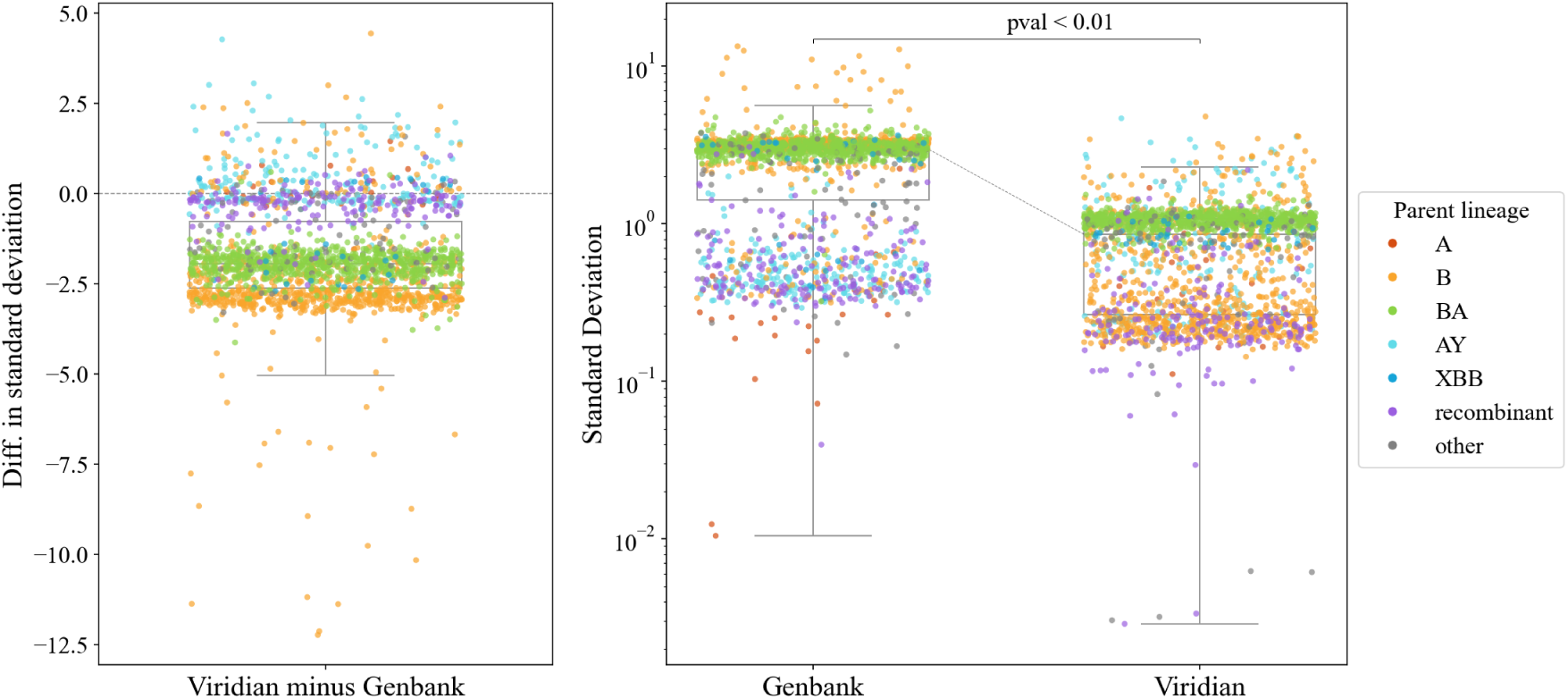
Comparison of uncertainty in growth estimates for different lineages when based on either the Viridian or Genbank tree. Panels a) (left) and b) (right) plot the same data in two ways; each point represents one lineage. Panel a) plots the difference in standard deviation of posterior density of relative growth rate estimate Δ log *R* (i.e. standard deviation using the Viridian tree minus standard deviation using the Genbank tree). Negative values here show that on average, the Viridian tree yields lower uncertainty than the Genbank tree. Panel b) shows the standard deviation of the posterior density of relative growth rate estimate Δ log *R* based on the GenBank tree (left) and Viridian tree (right). The median standard deviation of strain growth rate using the Genbank tree is 2.967, while the median standard deviation using the Viridian tree is 0.859. This difference is statistically significant (*p <* 0.01, paired t-test). Box-plots show first and third quartiles (lower and upper boundaries of box), and whiskers are set to the farthest point that is within 1.5 times the inter-quartile range from the box. Legend labels denote parent lineage.

### Final global tree and masking

We updated our global sample list to include data from the ENA/SRA as of 28^th^ June 2024, making a final global tree of the Viridian consensus sequences containing 4,471,579 samples. Tree construction was done, as is normal with UShER, by batching the samples, and then alternating adding a batch to the tree and optimising the tree. In the process of doing this, we noted how the order in which samples were passed to UShER had a very significant effect on the deep structure of the tree. Passing them in in random order resulted in the initial tree being constructed with recombinant genomes, resulting in considerable misplacement of the VOCs. We determined that the best approach was first to construct a tree with samples with no missing data, passed in in temporal order, then to add lower quality samples later (see Methods). After constructing the tree, we masked positions in the Problematic Sites set, which includes highly homoplasic sites in addition to sites previously observed to be reversion-prone in SARS-CoV-2, and masked 31 reversions that occurred 200 or more times in the tree – this choice of 200 allowed us to exclude position 11083 (highly homoplasic, and one of the first Problematic Sites), but did not include 23040 where there have been true reversions multiple times in Omicron. After masking, we ran matOptimize [26] to improve the structure of the tree in the absence of artefactual reversions and highly homoplasic sites.

### Impact on evolutionary and epidemiological analysis

The primary aim of this study is to provide a high quality resource (assemblies and phylogeny), with less “ad hoc masking”, with the intention that it reduces systematic error and noise in downstream work of others. We give two example applications.

First, in order to estimate the effect of the reduced number of sequence/assembly artefacts in the Viridian assemblies on epidemiological analysis, we used geographic metadata for each sample and a pandemic-scale cluster estimation algorithm (matUtils, Cluster-Tracker [27]), to compare the number of inferred unique SARS-CoV-2 viral introductions in each country using the GenBank and Viridian data (Supplementary Table S8). The expectation would be that removing artefactual errors would reduce the number of small clusters, caused by errors pushing genomes out of the larger clusters they truly belong in, creating artificial “introductions”. We found, for every country except Slovakia, there were more inferred introductions with the GenBank assemblies. The effect is more pronounced in highly sampled geographic regions, especially the United States (15,026 versus 13,626 introductions and 7,281 versus 6,676 singleton clusters for GenBank vs Viridian); see Supplementary Figure S13. As predicted, we see fewer small introductions with Viridian, and at the far right (note log scales) the very largest clusters are slightly larger.

Secondly, we quantified the extent to which the higher quality assemblies would affect estimates of differing mutational spectra of different Variants of Concern [28]. In all cases the spectra were very similar (i.e. the effect was limited), but interestingly in Alpha there had been an odd T*>*A context (labelled with an arrow in Supplementary Figure S14a) that was elevated above all others with the August 2022 UShER tree, which was gone in the Viridian data (Supplementary Figure S14b). The difference in G*>*T mutations that had been observed previously between Omicron and non-Omicron is still very much present; see Supplementary Figure S15 – confidence intervals (shown as error bars) do not always overlap the *x*=*y* line, so there are minor differences in the exact values but the overall trend and conclusions are unchanged.

## Discussion

The pandemic was met with an unprecedented globally-distributed sequencing effort that imposed substantial challenges for comparing and jointly analyzing data produced by thousands of labs with heterogeneous sampling, molecular, bioinformatic, and analysis protocols. In particular, the downstream effect of using multiple variable-quality genome assembly workflows, inconsistent QC criteria, and the inevitable co-evolution of virus and amplicon schemas, led to systematic errors in genomes, and therefore the phylogeny.

Here we present Viridian, a fast, low resource viral assembly tool specifically designed for tiled amplicon data and use it to produce a high quality sequence dataset of all publicly deposited SARS-CoV-2 data from January 2020 through to June 2024. With this we were able to build a much higher quality phylogenetic tree, needing less masking, than the current phylogeny.

We hope for three outcomes. First, that this resource will provide a valuable substrate for detailed methodological, evolutionary and epidemiological analyses. This has already happened, with de Maio et al developing new methods for handling mutation rate variation and sequencing errors in large phylogenies [29]. Second, that Viridian itself will prove useful, providing a significant improvement for Nanopore (and marginal for Illumina) compared with the ARTIC workflow, and a standardised single workflow and output format for Illumina, Nanopore and Ion torrent. Third, that in future epidemics or pandemics, the tools and ideas from this paper will serve to reduce the amount of time spent poring over trees and trying to distinguish artefact from biology. Viridian will work for tiled amplicon sequencing of non-segmented viruses where a consensus is the desired output (i.e. not in circumstances where multiple strains should be identified) and a single reference can be used. In other words, situations where there is limited structural variation or hypervariability, such as a particular outbreak, or a recent zoonosis (eg SARS-CoV-2).

We note that a similar approach (amplicon-by-amplicon assembly followed by remapping for QC) has been previously used for HIV (https://github.com/neherlab/hivwholeseq?tab=readme-ov-file#1-mappingfiltering-sample-by-sample). An alternative approach, more robust to handling hypervariable regions, is to do amplicon assembly followed by de novo scaffolding of amplicons without use of a reference. This method was implemented in the tool Lilo, used for African Swine Fever Virus [30].

Despite all this, bioinformatic methods can only go so far. Quality control within a single lab is relatively easy, especially if one can use molecular protocols, such as negative controls and using synthetic spike-ins [31]. However, maintaining quality levels from distributed sequencing and assembly on a national and global scale is much harder. Our approach (uniform reprocessing) is actually the simplest, providing the raw data remains available. However, it is not a viable approach mid-pandemic when there is barely enough time to keep up with incoming data. We therefore advocate for improved standardisation (and adoption) of metadata around sampling, assembly and QC, and also multinational “simulations” of pandemics to better prepare for integrating data from different pipelines.

Returning to our project, since the data in the ENA/SRA is heavily biased towards a few high income countries (especially USA and UK), we realised it was important to increase the geographical breadth of our dataset, and reached out to scientists around the globe inviting them to join our collaboration. This preprint represents Phase 1 of this project. Our team has now submitted pre-existing raw sequence data to the ENA/SRA from Argentina, Austria, Germany, Ghana, India, Netherlands, South Africa, Singapore and Sri Lanka. The worldwide distribution of samples is shown in Supplementary Figures S16,S17 (raw data is in Supplementary Table S9).

The pandemic was a global catastrophe with a huge cost in life, and the immense efforts of health professionals on the front lines, public health officials and the (sometimes ad hoc) networks of scientists and bioinformaticians has left many exhausted. However it is also a story of tremendous achievement and solidarity. In doing this work and building this collaboration, it has been striking how everyone has been determined to make the most of this vast resource of SARS-CoV-2 genomic data and build the cleanest and most correct assemblies and phylogeny as possible, to the benefit of us all. It has been a privilege to work together to produce these high quality resources, which was only possible because raw sequence data was deposited in the ENA/SRA.

## Methods

### Viridian pipeline

The main stages of the assembly process are: identify the amplicon scheme; sample the reads per amplicon; generate a consensus sequence by overlapping a consensus built for each amplicon; determine variants by aligning the consensus to the reference sequence; mask low quality bases using read mapping to the consensus, to output a final masked consensus sequence. An overview of the pipeline is shown in Figure 6.

**Figure 6:**
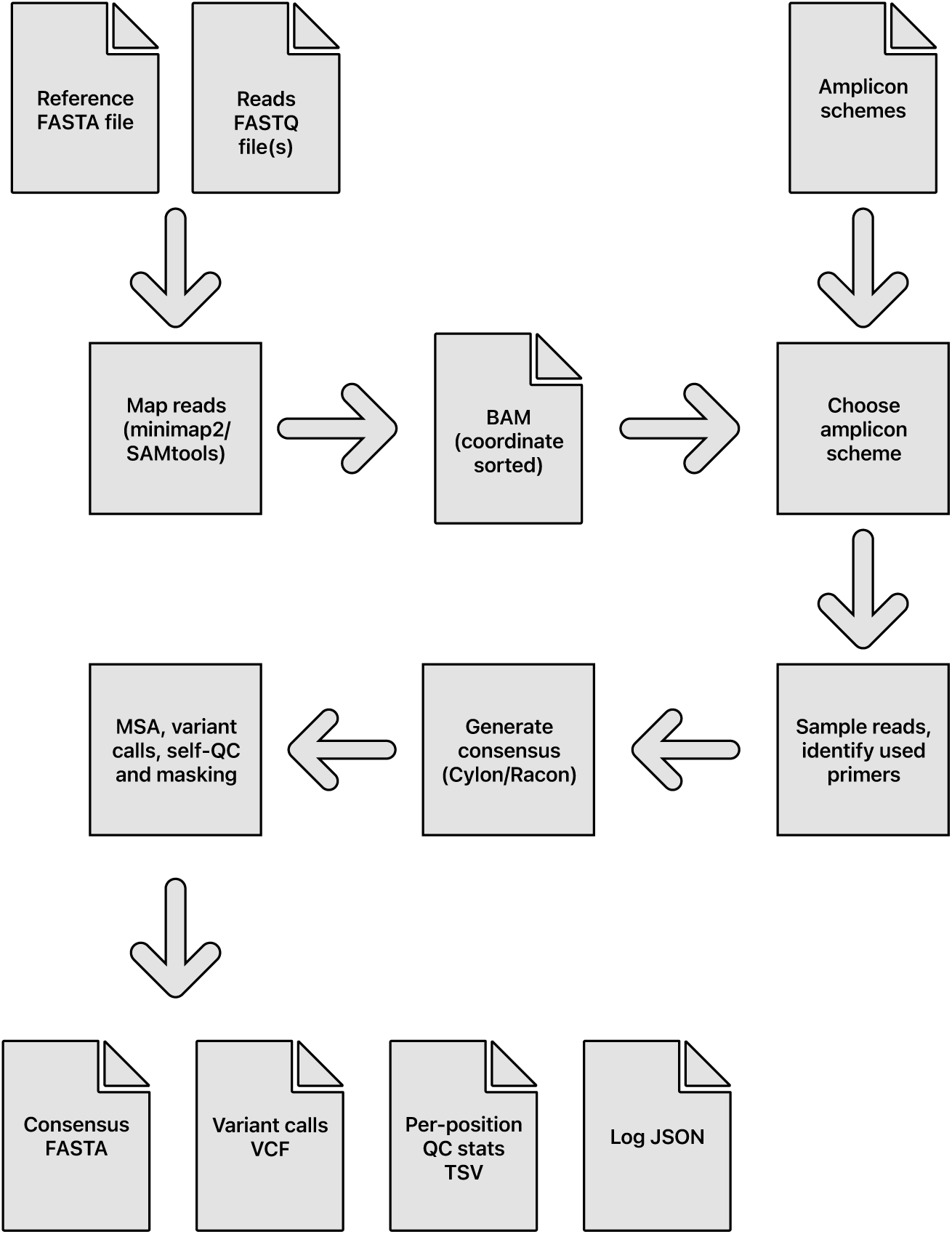
Overview of the Viridian pipeline, from input sequencing reads to output files.

#### Amplicon scheme identification

The amplicon scheme is automatically identified from the reads, from the built-in set of schemes (users can optionally add their own): AmpliSeq v1; ARTIC versions 3, 4.1, 5.3.2 400, 5.2.0 1200 [32]; Midnight 1200 [18]; and VarSkip v1a-2b (https://github.com/nebiolabs/VarSkip).

The reads are mapped to the reference genome (default SARS-CoV-2 MN908947.3) using minimap2 [33] with options –x map-ont (Nanopore) or –x sr (Illumina/Ion Torrent). SAMtools [34, 35] is used to make a sorted by coordinate and indexed BAM file, which by default is deleted at the end of the run but can be kept using the option –-keep bam. This BAM file is parsed using Pysam (https://github.com/pysam-developers/pysam) to determine read depth across the genome and which amplicon scheme is the best match to the reads. Mappings flagged as secondary or supplementary are ignored. If reads are paired then only proper read pairs are used. The pipeline is stopped at this stage if (by default) less than half of the genome has more than 20X read depth.

For each amplicon scheme under consideration, a normalised score is calculated based on the positions of mapped fragment ends. Throughout, “fragment” means the mapped portion of an unpaired read, or the leftmost to rightmost mapping coordinates of a proper read pair. The idea is that fragment end mapping positions are expected to stack up at the left end of left primers and the right end of right primers, since the reads are from amplicon sequencing. The score is an overall measure of how close the fragment ends are to the primer ends.

At each position in the genome, the number of fragments with leftmost mapped end at that position is counted. These counts are used to score each amplicon scheme separately in turn (Figure 7). For each position in the genome, the distance to the nearest left end of a left primer in the scheme is found, moving to the left of that position. For example, if there is a left primer at position 100-130, then (assuming no other primers in this region), position 103 would have a distance of 3 (see Figure 7(a)). Then at that position, we find how many fragments had their left end mapped at that position, and add that number to a counter of nearest distances. For example, if there were 20 fragments with left end at position 103, then 20 would be added to the counter for distance 3. The process is repeated for right primers, resulting in a count of mapped fragment ends at each distance from a primer (see Figure 7(b,c)).

**Figure 7:**
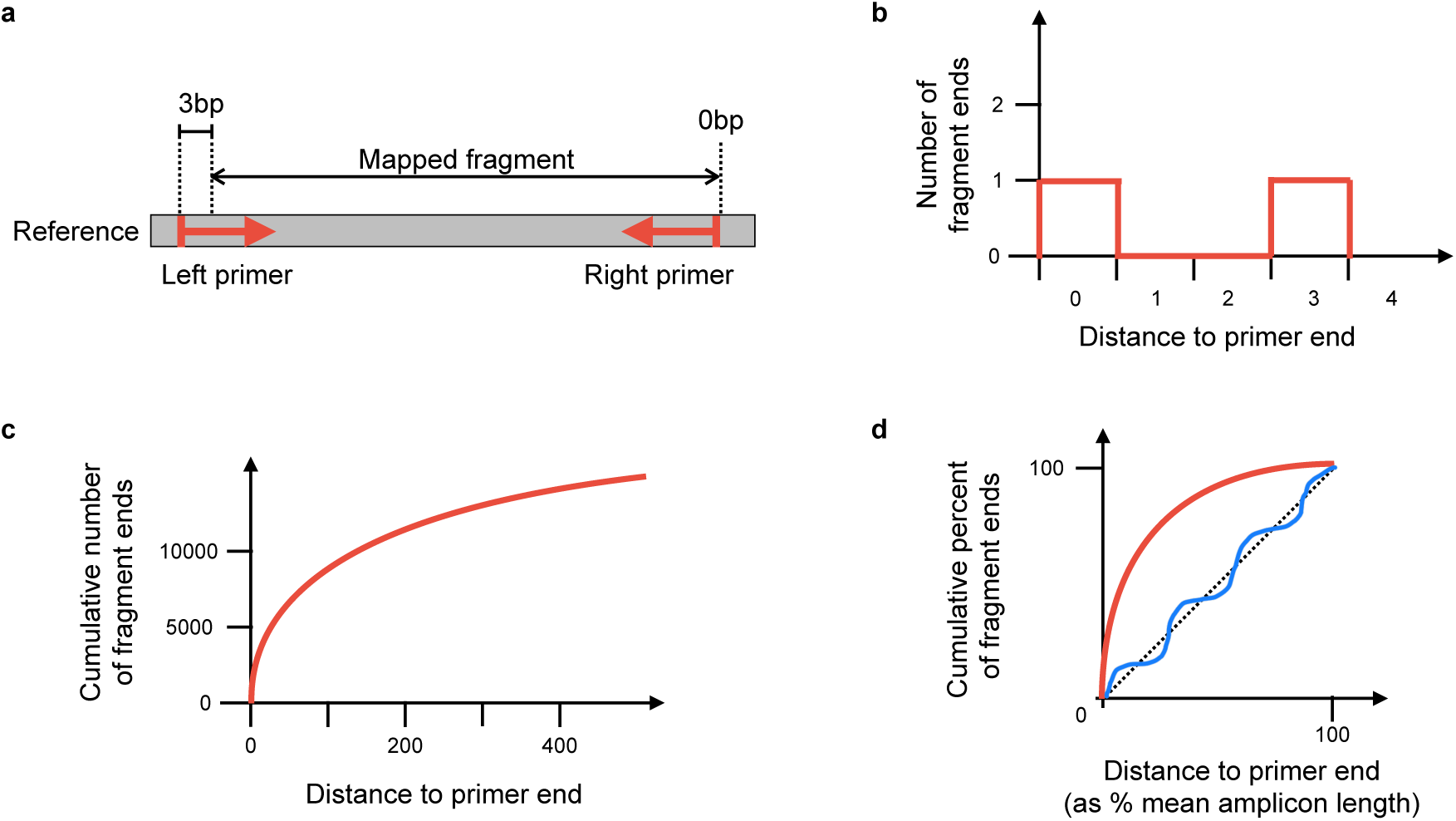
Method to score an amplicon scheme, using mapped fragments. **a)** Example of one mapped fragment, where its left end is 3bp from the start of the primer, and its right end is 0bp from the end of the right primer. **b)** The plot generated from the fragment in a). The right end of the fragment increments the counter for zero distance from a primer, and the left end of the fragment increments the counter for 3bp distance from a primer. The information from all fragments in the sample is added in this way, to make the distribution of distances from nearest primer ends. **c)** The cumulative plot from b) after adding all fragments. **d)** Plot c) is normalised by taking distance to primer end as a percentage of the mean amplicon length (*x* axis), and fragment counts as percent of total fragments (*y* axis). The red line indicates a typical curve where the reads match the scheme, whereas the blue line shows a scheme that does not match. The scheme’s score is the sum of differences between the calculated line and the *y* = *x* line (shown as a dashed line).

The distance is normalised by taking the distance as a percent of the mean amplicon length for the scheme, and the count of fragment ends is normalised by taking the percent of total fragment ends. The results are binned, so that for each integer *i* in the range 0 *−* 100, we know the percent of fragments *f* (*i*) ending normalised distance in the interval [*i, i* + 1) from a primer. The score is defined as

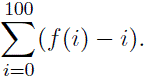

This is similar to calculating the area between the observed fragment counts and the line *y* = *x* (Figure 7(d)), but negative values are allowed. The maximum possible score for perfect reads is 5050, because f(i) = 100 for all *i* and the score is then

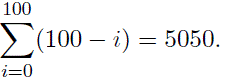

Intuitively, a scheme that matches the reads will have fragment ends close to the primer ends, resulting in an initial steep curve. Conversely, a scheme that is not related to the reads should approximately follow the line *y* = *x*. Therefore measuring the divergence from the *y* = *x* line provides a reliable measure of how well the scheme and reads agree. See Figure 7(d) for cartoons of a matching and non-matching scheme, and 8 for a real example output by Viridian. Viridian chooses the scheme with the highest score. However, if the best score is less than 250, or less than double the second best score, then the run is stopped and the sample is considered to be failed. For context, ERR8959196 shown in Figure 8 had best score 4290 and second best score 464.

**Figure 8:**
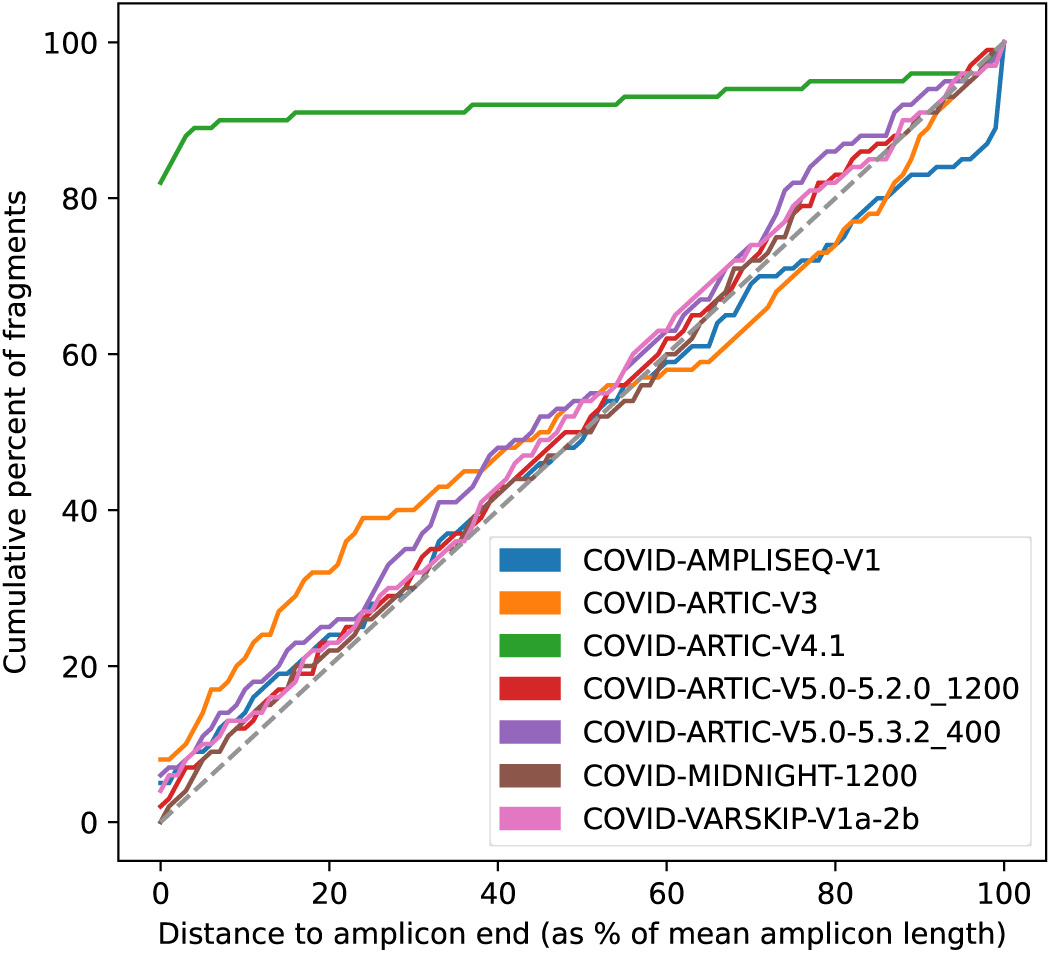
Example scheme identification score plot from Viridian. Made from run accession ERR8959196, which is Nanopore reads sequenced using ARTIC-V4.1 primers.

#### Read sampling

Once the amplicon scheme is known, reads are sampled to a target depth of (by default) 1000X for each amplicon, or using all reads for an amplicon if the mean depth is less than 1000X. If a fragment matches to more than one amplicon, then it is assigned randomly to one of the amplicons (the random number generator is seeded so that results are deterministic).

Within an amplicon, where there is more than one left primer (and similarly in the following description for right primers), the number of fragments supporting that primer is counted. Here, support is counted as the left fragment end being within 5bp of the start of the primer. A primer is excluded from the remainder of the pipeline if it is supported by fewer than 20 fragments. The exception is that if no left primers for the amplicon have support, then all left primers are kept. The result is an inferred amplicon scheme, consisting of a subset of the original primers from the chosen scheme.

Each fragment is assigned to a left and right primer pair within its designated amplicon. These are chosen by taking the rightmost left primer and leftmost right primer that contain the fragment. In summary, at this point in the pipeline we have a set of reads for each amplicon with mean coverage 1000X (or lower if there were not enough reads sequenced for an amplicon). Where an amplicon has more than one left and/or right primer, the set of reads is further split into sets for each primer pair.

#### Assembly

A consensus sequence is generated using a separate module called cylon (https://github.com/iqbal-lab-org/cylon). The overall method is to generate a consensus for each amplicon, overlap these consensus sequences into contigs, then scaffold against the reference sequence to output a final consensus sequence for the genome (Figure 9). It takes the inferred amplicon scheme (as described in the previous section) and a set of sampled reads for each amplicon. Reads are further sub-sampled for each amplicon from the 1000X reads, with a target depth of (by default) 150X for Illumina and 250X for Nanopore or Ion Torrent.

**Figure 9:**
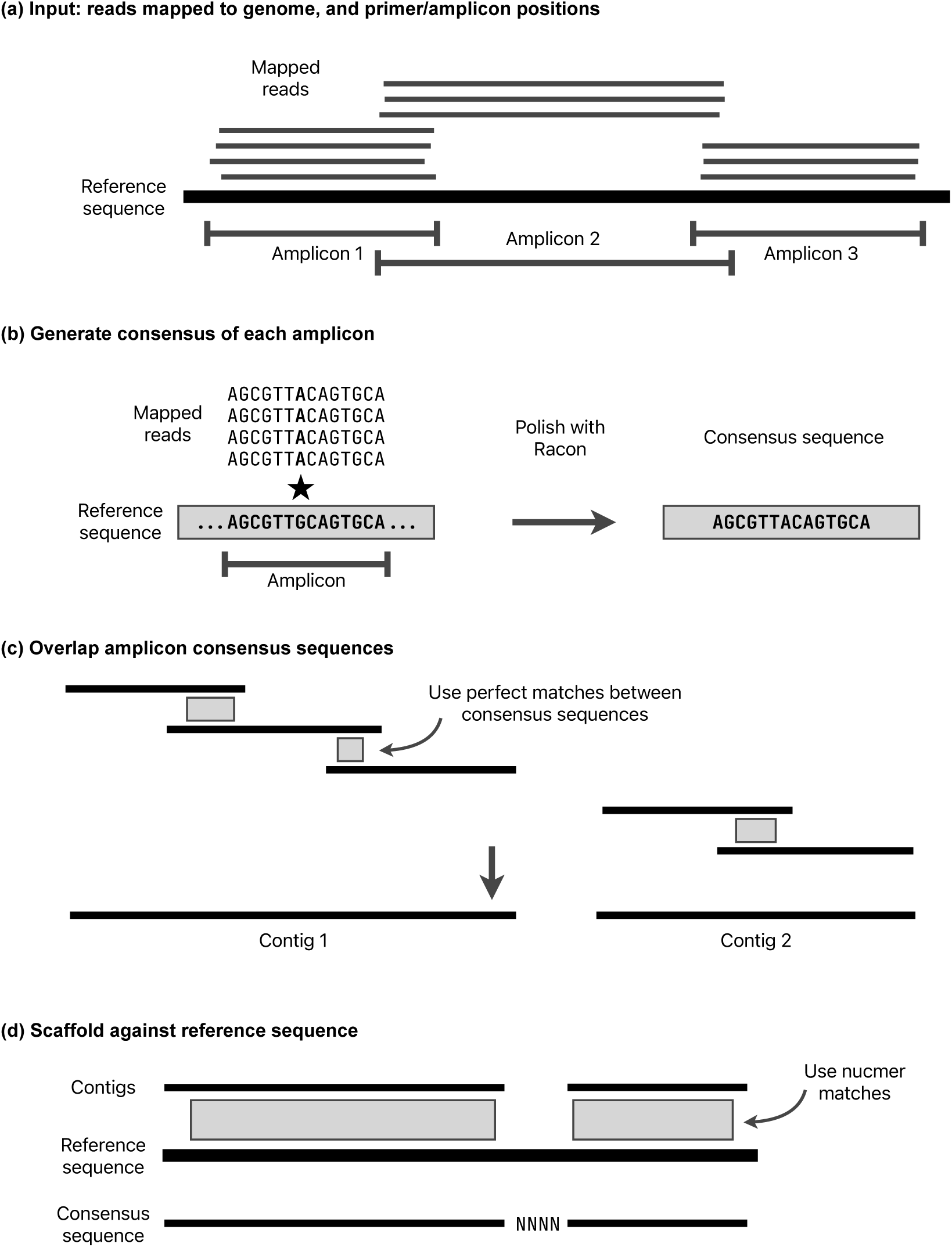
Consensus sequence construction methods. See main text for details. a) The starting point is primer and amplicon positions, and reads mapped to the consensus sequence. b) The consensus sequence of each amplicon is generated independently, using Racon. c) The amplicon sequences are overlapped using perfect matches (if they exist), making contigs. d) The contigs are scaffolded against the reference genome, adding gaps where needed.

A consensus sequence is generated for each amplicon by iteratively running Racon [20] until no more corrections are made, up to a maximum of 10 runs. If the input reads are paired, then each read pair is merged where possible using NGMerge [36] before running Racon. During testing, merging read pairs was found to improve the accuracy of Racon. In each Racon iteration, reads are mapped using minimap2 with options –x map-ont (Nanopore) or –x sr (Illumina/Ion Torrent). Racon options –-no-trimming –-window-length W are used, where W is the length of the amplicon plus 100 to avoid any erroneous indels at window ends. If no sequence is returned from Racon, then the amplicon is classed as failed. The sampled reads are mapped back to the consensus sequence and all positions with less than 5X depth are masked with Ns. If the resulting sequence is shorter than 30bp or has more than 50% Ns then the amplicon is failed.

Once there is a consensus sequence for each amplicon, adjacent amplicons are merged. First, amplicons are mapped to the reference genome using minimap2, and those with no mapping in the correct orientation are classified as failed and removed. If there is a perfect sequence match of at least 10bp between adjacent amplicons, it is used to join them. Otherwise, if the minimap2 match coordinates imply that adjacent amplicons overlap (the reference positions overlap), then those matches are used. Finally, if the minimap2 matches do not have overlapping reference positions, for example if one or both of the amplicons have a truncated consensus sequence, then a contig break is placed between the two amplicons.

Note that the start and end of the consensus sequence from each amplicon is excluded by this overlapping method, meaning that unreliable regions of consensus sequences that were inferred from reads starting or ending with primers are excluded. The only exception to this is where an amplicon is dropped, the next amplicon will include primer sequence. However, this is masked later in the QC stage. The amplicon overlapping is repeated for each adjacent pair of amplicons, stitching together a consensus sequence.

Once all possible adjacent amplicons have been merged, the result is one or more contig(s). When there is more than one contig, the position in the reference of each contig is determined using nucmer from the MUMmer software package [37]. The contigs are scaffolded, putting an estimated number of Ns between them based on the mapping coordinates. Since there could be insertions or deletions in the sample, this number of Ns is not reliable, but it is corrected during the next stage.

#### Variant calling

Variants are called with respect to the reference genome using the function make truth vcf from the tool varifier [38]. This globally aligns the cylon consensus sequence to the reference genome to identify variants. Since the amplicon schemes do not cover the complete reference genome, false-positive deletions are excluded from the start and end of the genome using the options –-global align min coord, –-global align max coord to restrict to coordinates within the amplicon scheme. Gaps in the consensus (ie strings of Ns) are corrected to be the same length as the corresponding portion of the reference sequence using the option –-sanitise truth gaps. These incorrect lengths can arise from failed amplicons, where the amplicon overlapping algorithm cannot always determine the exact gap length. For nanopore and Ion Torrent reads, indels of length 1 or 2 are removed from the consensus sequence using the option –-indel max fix length 2. This removes false-positive indels caused by the error model of those technologies, at the cost of excluding real calls. However, in most cases any true-positive call that is removed will be masked later in the QC and masking stage of the pipeline.

The end result of this stage is a VCF file of variants, a consensus sequence with consistent gap lengths, and the alignment of the reference and consensus sequences.

#### QC and masking

During read sampling to 1000X read depth per amplicon, each fragment (read pair or single unpaired read) is allocated to a left and right primer, by taking the smallest primer range that spans the entire fragment. For each amplicon and each primer pair within that amplicon, all reads for that primer pair are mapped to the consensus sequence using minimap2 (with the same options as the original run of minimap2) and then pileup is run to gather coverage statistics. Keeping the reads partitioned in this way means that at each genome position, the results from one pileup run can be counted as either inside a primer (“bad” coverage) or not inside a primer (“good” coverage). This is outlined in Figure 10. Pileup is calculated using the pileup function from pysam with the stepper option set to samtools, and ignore overlaps and compute baq set to False.

**Figure 10:**
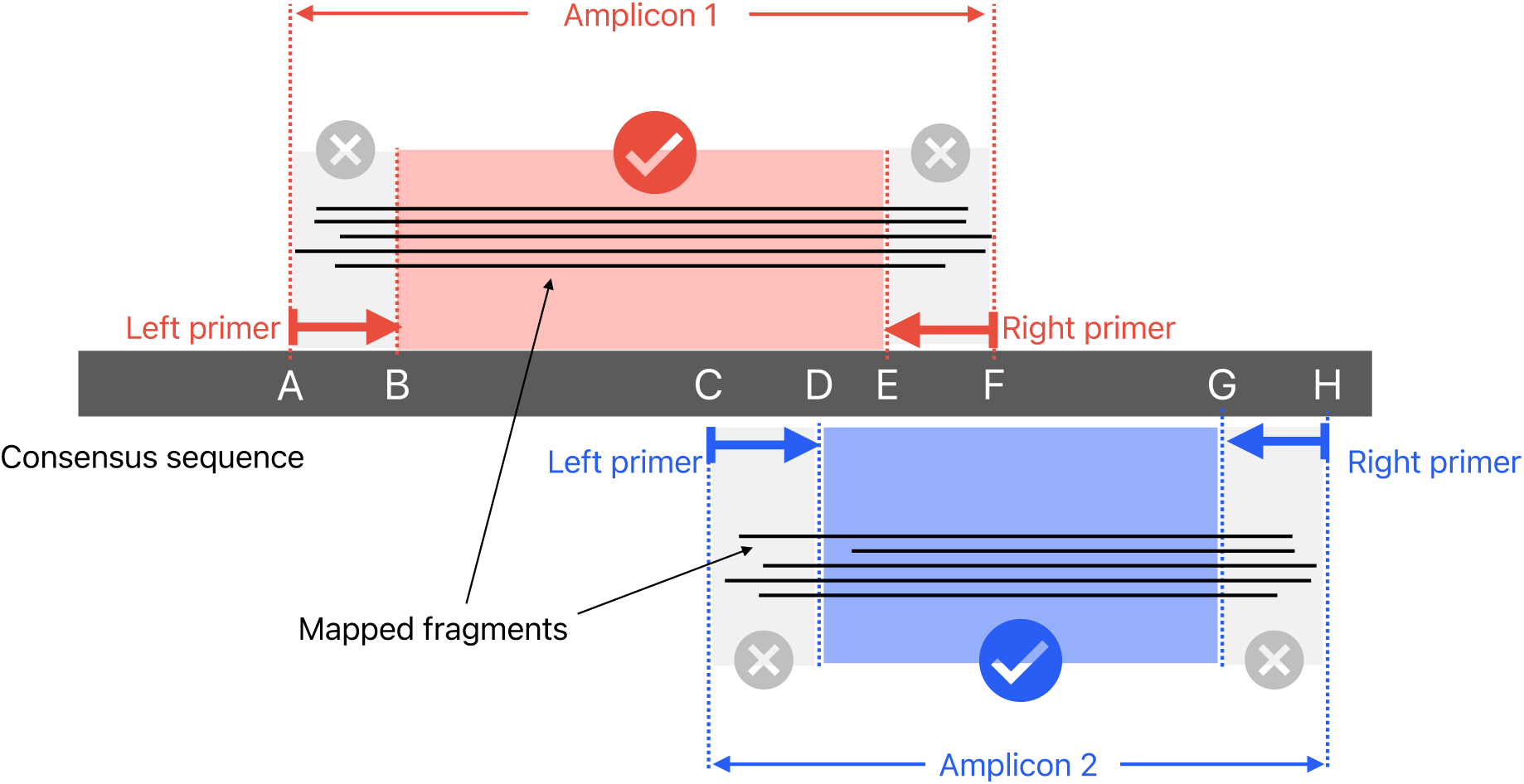
Consensus sequence pileup/masking methods. Two amplicons are shown with fragments (either illumina read pairs, or unpaired nanopore reads) mapped to the consensus. The fragments from amplicon 1 contribute to pileup at B-E, and do not count towards the primer regions A-B or E-F. Similarly, the fragments from amplicon 2 contribute to coverage at D-G (but not to C-D or G-H).

Pileup results are aggregated at each position in the consensus sequence. This is used with the reference genome/consensus sequence alignment to output a TAB-delimited report with read depth details at each position (split into separate counts for good and bad coverage). The good coverage is used to generate a masked consensus sequence, where untrustworthy positions are replaced with Ns. If the majority of reads disagree with the consensus position, or fewer than 20 reads in total agree with the consensus, then it is masked. At positions where there is evidence of more than one allele – by default an allele is counted as present if is supported by at least 20% of reads – then the consensus base is replaced with an ambiguous IUPAC code (for example, “R” to mean “A” or “G”).

#### Output files

The final masked consensus sequence is written in FASTA format, plus other files with additional information. Plots of read depth across the genome and scheme identification scoring are made. All QC results are written to a tab-delimited file with one position per row, including detailed read depth information. A log file in JSON format is written, with a high-level results summary section that includes all command line parameters, run time, version information and consensus sequence statistics. It also contains detailed information such as the MSA between the reference and consensus, amplicon details (chosen primers, number of matching reads etc), and genome-wide read depth statistics.

### Simulated data

We developed a Snakemake [39] pipeline to simulate PCR artefacts for 8000 SARS-CoV-2 samples, to compare the assembly accuracy of Viridian to the Connor Lab (https://github.com/connor-lab/ncov2019-artic-nf) and Epi2me labs (https://github.com/epi2me-labs/wf-artic) ARTIC Nextflow workflows. Firstly, truth assemblies are simulated from a reference genome and reference phylogeny using PhastSim [40] and truth variant calls obtained using varifier [38]. The primer sequences of the ARTIC v4 amplicon scheme are then mapped to the truth assembly of each sample using the aln command of bwa [41] to get the start and end positions of each amplicon and check for sequence mismatches in primer binding regions. If one or more mismatches are identified, one of two possible PCR artefacts are simulated with equal probability: either the primer sequence containing the mismatch is replaced with the reference sequence, or the amplicon is assigned a read depth of 0. Random amplicon dropout is simulated with probability 0.001 and the sequencing depth of all other amplicons is drawn from a Normal distribution (*µ* = 500*, SD* = 20). Reads are then simulated from each amplicon at the selected sequencing depths using ART [42] for Illumina and Badread [43] with –-identity 94,98.5,3 for Nanopore. The reads of each amplicon are aggregated such that there is one FASTQ of Illumina and one of Nanopore reads per sample and the reads are assembled using the Connor lab pipeline and Viridian workflow for Illumina and Epi2me labs pipeline and Viridian workflow for Nanopore. Finally, a new tool called Covid Truth Eval (https://github.com/iqbal-lab-org/covid-truth-eval), which is described in detail later, was used to generate TSV files that summarise the assembly accuracy for each tool.

### Empirical truth set

Combined nasal and oropharyngeal specimens were identified during routine sequencing at Oxford University Hospitals NHS Foundation Trust (OUH) as part of Pillar 1 national surveillance in the United Kingdom. Specimens were selected representing the Pango lineages B, B.1, B.1.1.7, B.1.1.7 (E484K), B.1.214.2, B.1.351, B.1.525, B.1.617.2, B.28, BA.1, P.1 and P.2. These were retrieved and cultured at the University of Oxford, generating abundant virus stocks. RNA from these virus stocks was sequenced using Illumina and Oxford Nanopore instruments with both ARTIC and ONT Midnight protocols, in addition to sequence-independent single-primer amplification, forming the dataset deposited in ENA projects PRJEB50520 and PRJEB51850 [44]. Sequencing was performed at the University of Oxford except where otherwise stated below.

#### Viral culture

Vero cells were maintained in Dulbecco’s Modified Eagle Medium (DMEM) high glucose supplemented with 1% fetal bovine serum, 2mM Glutamax, 100 IU/ml penicillin-streptomycin, and 2.5*µ*g/ml amphotericin B at 37°C, 5% CO2 in a humidified atmosphere before inoculation with 200*µ*l of throat swab fluid. Cells were then incubated at 37°C, with daily monitoring for cytopathic effects (CPE). When CPE reached 80%, virus-containing supernatants were harvested through centrifugation at 3,000 rpm at 4°C and stored at –80°C in single-use aliquots. Virus titers were quantified by a focus-forming assay on Vero cells. Spike genes were sequenced in order to verify protein sequence integrity. Refer to [45] for more details.

#### Extraction

Viral RNA was extracted from 200*µ*l and 400*µ*l volumes of Coplan viral transport media on the KingFisher Flex system (Thermo Fisher, UK) using the MagMAX Viral/Pathogen II Nucleic Acid Isolation Kit (IVD). Two wash steps were incorporated and extracts were eluted in 50*µ*l.

#### PCR

PCR tests were performed by OUH using two PCR assays: Altona RealStar (targeting E and S genes; Altona Diagnostics, Liverpool, UK) and Thermo Fisher TaqPath assay (targeting S and N genes, and ORF1ab; Thermo Fisher, Abingdon, UK).

#### Sequence Independent Single Primer Amplification (SISPA)

Viral RNA was extracted as described above then Complementary DNA (cDNA) was prepared using a SISPA approach [46]. In brief, firstly RNA was reverse-transcribed with SuperScript III Reverse Transcriptase (Life Technologies, UK) using Sol-Primer A (5’-GTTTCCCACTGGAGGATA-N9-3’)

[47]. Then 5*µ*L of cDNA and 1*µ*L (100pmol/*µ*L) Primer B (5’-GTTTCCCACTGGAGGATA-3’) were added to a 50*µ*L reaction using AccuTaq LA (Sigma, Poole, United Kingdom), according to manufacturer’s instructions. PCR conditions were 98°C for 30s, followed by 30 cycles of 94°C for 15s, 50°C for 20 s, and 68°C for 5 min, and a final step of 68°C for 10 min. Amplified cDNA was purified using a 1:1 ratio of AMPure XP beads (Beckman Coulter, Brea, California, US) and quantified using the Qubit High Sensitivity dsDNA kit (Thermo Fisher Scientific, UK).

#### SISPA Oxford Nanopore sequencing

SISPA products were sequenced following a previously described protocol [48] using Oxford Nanopore Technologies (ONT) native barcoding (EXP-NBD104) and ligation sequencing (SQK-LSK109) kits with R9.4.1 flow cells.

#### ARTIC V3 Illumina sequencing

Libraries were prepared using the NEBNext^®^ ARTIC SARS-CoV-2 Library Prep Kit, following standard protocol with cDNA Amplicon and Ligation Bead Clean-ups (Version 3.0 7/21). Manual library normalisation was performed to ensure even sample coverage, based on the library’s DNA concentration and average size, as measured by the Qubit (Thermo Fisher Scientific, UK) and 2200 TapeStation (Agilent Technologies, USA). Paired-end sequencing was performed using the MiSeq reagent kit v2, with 2×250bp, and one water control on each run. NEBNext^®^ Multiplex Oligos for Illumina^®^ (96 Unique Dual Index Primer Pairs) were used.

#### ARTIC V4.1 Illumina sequencing

Libraries were sequenced at the University of Northumbria following the ARTIC V4.1 CoronaHiT-Illumina protocol [49], using an Illumina NextSeq 550.

#### ARTIC V3 Oxford Nanopore sequencing

Sequencing was performed using the ARTIC LoCost protocol and v3 primers using R9.4.1 flow cells. Final library concentration was quantified by the High Sensitivity dsDNA kit Qubit (Thermo Fisher Scientific, UK).

#### ONT Midnight Oxford Nanopore sequencing

Libraries were prepared using ONT Midnight RT-PCR Expansion kits (EXP-MRT001) and rapid barcoding (SQK-RBK110.96), following manufacturer protocols. R9.4.1 flow cells were used.

#### Manual curation

All reads were mapped to the reference genome MN908947.3 using minimap2 with the –x preset map-ont for Nanopore reads and sr for Illumina. A sorted BAM file was made using samtools sort. This was used to make an unfiltered set of variant calls by piping the output of samtools mpileup into bcftools call –vm. Each sample was curated manually, using Artemis [50] to view the mapped reads and infer a truth set of variant calls. Although the unfiltered calls from bcftools were used as a guide, the whole genome for every sample was inspected for variant calls. In rare cases where the Nanopore and Illumina reads disagreed at a position, it was flagged as “unknown”. The VCF files and metadata are available at https://github.com/iqbal-lab-org/covid-truth-datasets.

### Consensus accuracy evaluation

The accuracy of results of the simulated data and truth set were evaluated using a new tool CTE (Covid Truth Eval). It can evaluate either a VCF file of variant calls, or a consensus sequence, by comparing it with a “truth” consensus sequence. If the input is a VCF file, the consensus sequence to be evaluated is made by applying the variants to the reference sequence. It makes a multiple sequence alignment (MSA) of the consensus, truth, and reference sequences using MAFFT [24]. Each position in the genome is classified by comparing the base calls of the MSA, to verify the accuracy of the consensus sequence. The most common case is that the truth nucleotide is equal to the reference nucleotide, and the consensus also called the reference nucleotide. The possibilities for the truth are: a reference call, “homozygous” SNP (ie A, C, G, T that is different from the reference), “heterozygous” SNP (ie a mix of A, C, G, T), indel, dropped amplicon, or an N. Although rare, an N is used when the truth is unknown, as described above in the manual curation section. The possibilities for the consensus call are the same, except each nucleotide call could be correct or incorrect (ie the same as or different from the truth nucleotide). CTE reports the total count of each combination seen in the input sample.

Dropped amplicons are known in the truth data. However, they must be estimated from the consensus sequence that is under evaluation. Since tools can use different methods to mask a nucleotide or an entire amplicon, defining a position with an N as part of a dropped amplicon, or simply masked, is ambiguous. CTE uses the minimum possible range of coordinates we would expect to be Ns if an amplicon is dropped, ranging from one past the end of the previous amplicon to the position before the start of the next amplicon. If a run of Ns contains this range of coordinates for a given amplicon, then it is considered as dropped in the sequence under evaluation. Hence there is some ambiguity between “called as N” and “dropped” when interpreting the output of CTE.

### Africa dataset

The Africa dataset comprises a total of 12,287 samples, each of which has a “GISAID” assembly, and either Illumina (*N* =9935) or ONT (*N* =2352) sequencing reads, with primer schemes ARTIC Version 3 or 4, or MIDNIGHT-1200 (Supplementary table S7). All samples were processed with Viridian and ARTIC-ILM/ONT, producing a consensus sequence.

### Global dataset

Metadata for all sequencing runs with taxon ID 2697049 was downloaded using the ENA portal query https://www.ebi.ac.uk/ena/portal/api/search?result=read_run&query=tax_id=2697049&fields=all&limit=10000000 on 2^nd^ March 2023. These runs were filtered to only keep those with library strategy equal to AMPLICON, library source equal to VIRAL RNA, host empty or equal to homo sapiens, and instrument platform one of ILLUMINA, OXFORD NANOPORE or ION TORRENT. The resulting 5,288,952 sequencing runs were downloaded using either prefetch/fasterq-dump from the SRA-toolkit (https://github.com/ncbi/sra-tools) or enaDataGet (https://github.com/enasequence/enaBrowserTools). They were processed with Viridian, with 4,395,655 passing its QC requirements and producing a consensus sequence. These were further filtered for quality, requiring no more than 3 “heterozygous” base calls (ie none of A,C,G,T,N) and no more than 5,000 Ns. The N count was taken from the consensus sequence after aligning to the reference using MAFFT, as described in the Trees section later. A further round of filtering was applied based on requiring a reliable date for each sequencing run, using where available the collection date from the ENA/SRA, COVID-19 Genomics UK Consortium (COG-UK), and GISAID. Runs with no collection date from any source were removed. Where dates conflicted for a given sample, the order of preference used was the date with highest resolution, then COG-UK, GISAID, and finally ENA/SRA. At this stage, there were 3,960,704 runs – this is the set of runs used to compare with GenBank sequences (see next paragraph). Finally, the data were updated on 28^th^ June 2024, adding all new runs that passed the same QC requirements, making a total of 4,484,157 consensus sequences.

All GenBank genomes were downloaded on 23^rd^ May 2023 using the Datasets tool (https://github.com/ncbi/datasets) with parameters download virus genome taxon SARS-CoV-2. The genome and metadata files (genomic.fna.gz, data report.jsonl.gz) were extracted from the downloaded zip file. Genomes with host taxon ID (“host”*→* “taxId”) 9606, ie human, were kept. The genomes were matched to sequencing runs from the ENA/SRA using the run accession. Only GenBank genomes that matched to a single run that also belonged to the set of 3,960,704 Viridian consensus sequences (from the initial data obtained on ^nd^ March 2023) were kept. This resulted in an “intersection set” of 3,006,407 runs with both a Viridian consensus sequence and GenBank genome.

### Primer scheme validation

Since the COG-UK metadata includes the ARTIC primer scheme version, we used their project PRJEB37886 (included in the global dataset) to validate the scheme calls from Viridian. The ARTIC primer scheme version used was obtained from the SRA metadata using efetch (https://www.ncbi.nlm.nih.gov/books/NBK179288/) to download metadata for experiments in batches using the options –format xml –db sra –input ids.txt, where ids.txt is the name of the file containing a list of experiment accessions. The primer scheme version was extracted for each experiment from the value of the artic primer version tag in the EXPERIMENT ATTRIBUTES section of the XML data. Each efetch command was attempted twice (failures were common), resulting in a total of 2,485,169 primer scheme calls from ENA/SRA metadata. We then restricted to Illumina and Nanopore samples that passed Viridian (the 4,395,655 samples described earlier), and only included ENA/SRA primer scheme values of 3/ARTIC v3 for ARTIC version 3 and 4/4.1alt/ARTIC v4 for ARTIC version 4. This was a total of 2,341,118 samples.

Discordant samples for manual inspection were chosen by taking all Illumina samples with ENA/SRA scheme version 3 and Viridian scheme version 4, sorting by run accession, and taking 5 equally spaced runs from the list. The same method was used for Illumina with ENA/SRA version 4 and Viridian version 3, and then similarly for Oxford Nanopore samples, totalling 20 samples for manual inspection. Reads were mapped using minimap2 with the option –a to make SAM output, and the preset –x of sr (Illumina) or map-ont (Nanopore). A sorted BAM file was made using SAMtools, and then manually inspected with Artemis.

### Trees

Trees were built using MAFFT and UShER [22] and visualised with taxonium [51]. Each sequence was aligned to the reference using MAFFT with the option –-keeplength to force the alignment to be the same length as the reference genome, by only allowing gaps in the query sequence. The alignment was modified by forcing any gaps in the query sequence to be the same as the reference sequence. The resulting sequences were batched into sets of size 100,000. A VCF file was made for each batch with faToVcf, with the option –includeNoAltN. A tree was built by adding each batch in turn using usher-sampled and the option –-sort-before-placement-3. The final tree was optimized with the UShER command matOptimize and the options –m 0.000000001 –r 8 –T 20. Finally, the taxonium input file was generated using the script usher to taxonium from taxoniumtools [51]. The processing of input sequences to obtain taxonium input was implemented in a pipeline called Ushonium (https://github.com/martinghunt/ushonium).

In order to maintain an accurate tree structure, we ordered the samples by first using the samples with zero N or heterozygous calls, sorted by collection date. Then the remaining samples were used, again sorted by collection date. An exception to the date ordering was the 12,953 samples (3,876 of these were in the intersection set of 3,006,407 samples) where the GISAID date was given priority over other sources, which were added at the end instead of using the date. Using the highest quality consensus sequences first meant that UShER did not have to impute any ambiguous positions in any sequences. Sorting in date order meant that recombinant genomes – which emerged later in the pandemic – were not added to the tree too early, since they could be placed in an incorrect clade and then cause structural errors.

The global Viridian tree was built in two stages. A first version of the tree was built from the runs up to the 2^nd^ March 2023, using the order described above (highest quality and earliest collection date first). Positions in the Problematic Sites set (https://github.com/W-L/ProblematicSites_SARS-CoV2) were masked globally in the tree, and 31 reversions found to occur at least 200 times in the tree were also masked globally (all masked positions are listed in Supplementary Table S10). matOptimize was run following the masking to join branches that had been split by the masked substitutions or reversions. This tree was used as a starting point to update using the second batch of data from 28^th^ June 2024, with the same ordering method. The Problematic Sites positions were masked in new sequences before they were added to the tree. After the new sequences were added, in addition to masking the 31 reversions that occurred at least 200 times in the first version of the tree prior to masking, we added branch-specific masking for regions in BA.1 and BA.2.86 in which mafft misinterprets a deletion and insertion in close proximity as a series of substitutions. Positions 6513, 6515, 22195, 22197-8, 22202 and 22204 were masked in the BA.1 branch. Positions 21610, 21612-3, 21615-7, 21619-21, 21624-7, 21629, 21632, 21637, and 21639-41 were masked in BA.2.86. matOptimize was run after masking. 12,578 duplicate runs were removed from the tree that came from shared samples, to make a final tree with 4,471,579 unique samples/runs. We note that there are only 14 duplicate runs in the intersection tree, which were not removed.

### Py*R*_0_ analysis

Py*R*_0_ was run using Python version 3.10. Code is available online via GitHub at https://github.com/broadinstitute/pyro-cov?tab=readme-ov-file.

Analysis was done using the matched Viridian tree and Genbank tree of the intersection dataset. Py*R*_0_ estimates growth rate of lineages using a hierarchical regression model (see [25] for details); based on this, standard deviation of strain growth rate was aggregated across regions (countries, or first-level country divisions — e.g., state, province — if the first-level division has at least 50 samples) by summing region-specific standard deviations. A paired t-test was conducted on the standard deviation in growth rate estimates using the Viridian tree vs. Genbank tree. Supplementary Manhattan plots (spike protein and whole genome) only show mutations that appeared in both Viridian and Genbank trees, and a paired t-test was conducted on the growth rate estimates for each mutation. An unpaired t-test was also conducted on the full set of mutations, including those that only appear in the Viridian or Genbank trees, though no statistically significant results were found. Accompanying each Manhattan plot (Supplementary Figures S10,11) is a plot of the ratio of growth-related mutations to all mutations, where growth related mutations are defined as those which are at least one standard deviation from zero. Fisher’s Exact test was performed to analyze the difference in proportions of growth-related mutations in each annotated subdomain/reading frame of the spike protein/whole genome (respectively). To produce Supplementary Figure S12, rank was assigned according to the mean of the posterior density of the relative growth rate of a strain compared to the ancestral strain (denoted by *R/RA*) divided by the standard deviation of said posterior. Δ*logR* is the common log of the *R/RA* growth rate estimate. Mutation relative growth rate describes the relative growth rate conferred by a mutation compared to no mutation.

### Calculation of mutational spectra and proportions of G***>***T mutations

Mutational spectra were calculated as reported previously [28]. Briefly, all mutations downstream of the corresponding lineage root node are identified. The contexts of these mutations are calculated in the genomic sequence at the start of the corresponding phylogenetic branch, i.e. taking into account mutations that have arisen on ancestral branches in the phylogenetic tree. Mutational spectra were rescaled by the genomic composition in the lineage root ancestor as described previously [28]. Confidence intervals on the proportion of G*>*T mutations were calculated using Wilson score interval incorporating the calculated proportion and the number of sampled mutations.

### Software versions

Package versions used for the simulations were: Snakemake v7.8.5 [39], PhastSim v0.0.4 [40], ART v2016.06.05 [42], Badread git commit c2bdcbe [43], ARTIC Illumina workflow git commit 8af5152 from https://github.com/connor-lab/ncov2019-artic-nf, Epi2me wf-artic git commit 218aa1d from https://github.com/epi2me-labs/wf-artic, CTE git commit 9cd94b8 from https://github.com/iqbal-lab-org/covid-truth-eval, Nextflow v21.04.3 [52], bwa git commit c77ace7 [41], htslib v1.14 [53], samtools v1.14 [34], BEDTools v2.30.0 [54], joblib v1.1.0 from https://github.com/joblib/joblib, numpy v1.22.1 [55], pandas v1.4.0 [56], pysam v0.18.0 at https://github.com/pysam-developers/pysam and tqdm v4.62.3 from https://github.com/tqdm/tqdm.

The ARTIC-ILM pipeline used was git commit 8af5152 from https://github.com/connor-lab/ncov2019-artic-nf. The ARTIC-ONT pipeline used was git commit 218aa1d from https://github.com/epi2me-labs/wf-artic. Version 4.3 of Pangolin, and version 1.21 of pangolin-data were used for the intersection dataset. Version 1.29 of pangolin-data was used on the final Viridian global tree.

Viridian v1.0.0 or v1.1.0 was used to process all runs. The only difference between these versions is v1.1.0 added support for unpaired Illumina reads. The versions of tools used by Viridian were: Cylon git commit 57d559a, minimap2 git commit b0b199f, MUMmer v4.0.0rc1, NGMerge git commit 224fc6a, Racon git commit a2cfcac, Varifier git commit 8bc8726. Ushonium git commit b024320 was used, with dependencies MAFFT v7.520, UShER git commit 2df81ee, and taxoniumtools v2.0.111.

## Code and Data availability

Viridian is freely available under the MIT license at https://github.com/iqbal-lab-org/viridian. Supplementary text and figures S1-9 are in the supplementary PDF file. Code used for analysis and to generate figures is availble at https://github.com/martinghunt/viridian-paper. The global Viridian tree is hosted at https://viridian.taxonium.org. All other additional files are available from Figshare:

- Supplementary table S1, https://doi.org/10.6084/m9.figshare.27195261 – this is a TSV file containing metadata of all 5,959,032 sequencing runs considered in this study
- Supplementary tables S2-10 in one xlsx file, https://doi.org/10.6084/m9.figshare. 27195315.v2:
  S2 – Summary of counts of amplicon schemes in INSDC metadata and the scheme called by Viridian
  S3 – Accuracy of Viridian, ARTIC-ILM and ARTIC-ONT on simulated data
  S4 – Accuracy of Viridian, ARTIC-ILM and ARTIC-ONT on Illumina truth data set
  S5 – Accuracy of Viridian, ARTIC-ILM and ARTIC-ONT on Nanopore truth data set
  S6 – Run times and RAM usage on the truth data set
  S7 – Metadata for the African data set
  S8 – Numbers of inferred viral introductions
  S9 – Country counts in the Viridian global tree, and number of new samples since the tree was built
  S10 – Positions masked when building the global Viridian tree
- Supplementary HTML file, https://doi.org/10.6084/m9.figshare.25713198 – comparison of Viridian and GenBank assemblies
- All Viridian consensus sequences that are in the global tree, split over two tar archive files (https://doi.org/10.6084/m9.figshare.25713225, https://doi.org/10.6084/m9.figshare.27194637), which contain the sequences split over multiple xzipped FASTA files. These are the same batched FASTA files used when building the trees.
- The Viridian global tree of 4,471,579 sequences, in JSONL and.pb format, https://doi.org/10.6084/m9.figshare.27194547
- The GenBank and Viridian intersection trees in JSONL and.pb format, https://doi.org/10.6084/m9.figshare.25713285
- All other Viridian consensus sequences that are not in the global tree, split over two zxipped FASTA files https://doi.org/10.6084/m9.figshare.25713342, https://doi.org/10.6084/m9.figshare.27194652.

## Author contributions

Martin Hunt wrote the final implementation of Viridian, developed the primer-scheme identification system, assembled the genomes, developed the pipeline for tree-building and performed all analyses not listed below. Angie Hinrichs analysed and quality controlled the phylogenies and wrangled metadata. Daniel Anderson developed the simulation framework and performed analyses. Lily Karim performed the reversion analyses and geographical/introduction analysis. Bethany Dearlove analysed the assemblies, Ns, indels, and Pango assignments. Jeff Knaggs contributed to the first implementation of Viridian and initial exploratory work. Ben Kötzen performed the pyR0 analyis, with supervision from Jacob Lemieux. Piyada Supasa, Wanwisa Dejnirattisai, Chang Liu, Juthathip Mongkolsapaya and Gavin R. Screaton isolated and cultured virus stock used to construct the empirical truth set. Hermione Webster, Gillian Rodger, Teresa Street and Sheila Lumley sequenced the empirical truth set, and Bede Constantinides analysed and quality controlled the data. Philip Fowler did independent testing on different simulations and empirical data. Martin Hunt, Jeff Knaggs, Bede Constantinides, Tim Peto, Derrick Crook analysed the empirical truth set results. Theo Sanderson integrated the phylogeny into taxonium. Nicola de Maio did extensive quality control analyses of the genomes. Christopher Ruis performed the mutation spectrum analysis. Houriiyah Tegally, San Emmanuel James, Tulio de Oliveira collated the “early Omicron” dataset. All other authors collected samples, sequenced genomes and shared data with the archives. Zamin Iqbal conceived of the project. Zamin Iqbal and Russell Corbett-Detig supervised the project. Martin Hunt, Angie Hinrichs, Lily Karim, Daniel Anderson, Theo Sanderson, Russell Corbett-Detig and Zamin Iqbal wrote the paper. All authors reviewed the manuscript.

## Acknowledgements

We would like to thank Emma Hodcroft, Richard Neher, Duncan MacCannell, Nick Goldman, Trevor Bedford, Rachel Colquhoun, Andrew Rambaut, Robert Davies, and Rob Lanfear for discussions and advice. We also thank Zahra Waheed, Nadim Rahman and Khadim Gueye for help with ENA submissions. We thank the Global Pathogen Analysis Service team in Oxford, who used an early version (v0.3.7) of Viridian in production for over a year, providing valuable feedback. We thank the microbiology laboratory staff of the John Radcliffe Hospital, Oxford University Hospitals NHS Trust, for providing assistance with sample processing. We thank the IMSSC2 Laboratory Network Consortium members at the Robert Koch Institute for providing raw data sequences, the Sequencing Core Facility of the Genome Competence Center (MF1) at Robert Koch Institute for providing excellent sequencing services, and we thank all labs contributing to the German SARS-CoV-2 surveillance. We acknowledge high-performance computing services provided by the Robert Koch Institute. We gratefully acknowledge all data contributors, i.e., the Authors and their Originating laboratories responsible for obtaining the specimens, and their Submitting laboratories for generating the genetic sequence and metadata and sharing via the GISAID Initiative. Finally, we would like to thank Nick Goldman, who gave us the idea for the start of the introduction.

