## Supplementary Material for "Addressing pandemic-wide systematic errors in the SARS-CoV-2 phylogeny"

IMSSC2 Laboratory Network Consortium:
Barbara Biere, Ralf Dürrwald, Christin Mache, Djin-Ye Oh, Jessica Schulze, Marianne Wedde, Thorsten Wolff, Unit "Influenza and other Respiratory Viruses", RKI, Berlin, Germany; Stephan Fuchs, Torsten Semmler, Sofia Paraskevopoulou, Unit "Genome Competence Centre", RKI, Berlin, Germany; Romy Kerber, Stefan Kröger, Walter Haas, Unit "Respiratory Infections", RKI, Berlin, Germany; Konrad Bode, MVZ Labor Dr Limbach, Heidelberg, Germany; Victor Corman, Institute of Virology, Charité-University Medicine, Berlin, Germany; Michael Erren, MVZ Laborzentrum Weser-Ems; Patrick Finzer, MVZ Düsseldorf-Centrum, Düsseldorf, Germany; Roger Grosser, Labor Dr Wisplinghoff, Köln; Manuel Haffner, MVZ Labor Dr Kirkamm, Mainz, Germany; Beate Hermann, MVZ Dianovis, Greiz, Germany; Christina Kiel, MVZ Labor Dessau, Dessau-Roßlau, Germany; Andi Krumbholz, Thomas Lorentz, Labor Dr Krause, Kiel, Germany; Kristian Meinck, IMD-Laborverbund, Greifswald, Germany; Andreas Nitsche, Unit "Highly pathogenic viruses", RKI, Berlin, Germany; Markus Petzold, Institut für Medizinische Mikrobiologie und Hygiene, Institut für Virologie, TU Dresden, Germany; Thomas Schwanz, Institut für Medizinische Mikrobiologie und Hygiene, Universitätsmedizin Mainz, Germany; Florian Szabados, Laborarztpraxis Osnabrück, Georgsmarienhütte, Germany; Friedemann Tewald, Labor Enders, Stuttgart, Germany; Carsten Tiemann, Labor Krone, Bad Salzuflen, Germany.

#### Pandemic timeline

The same timeline as in the main Figure 1 is shown in Supplementary Figure S1, but with plots added showing the number of masked sites and nodes in the global phylogenetic tree of SARS-CoV-2.

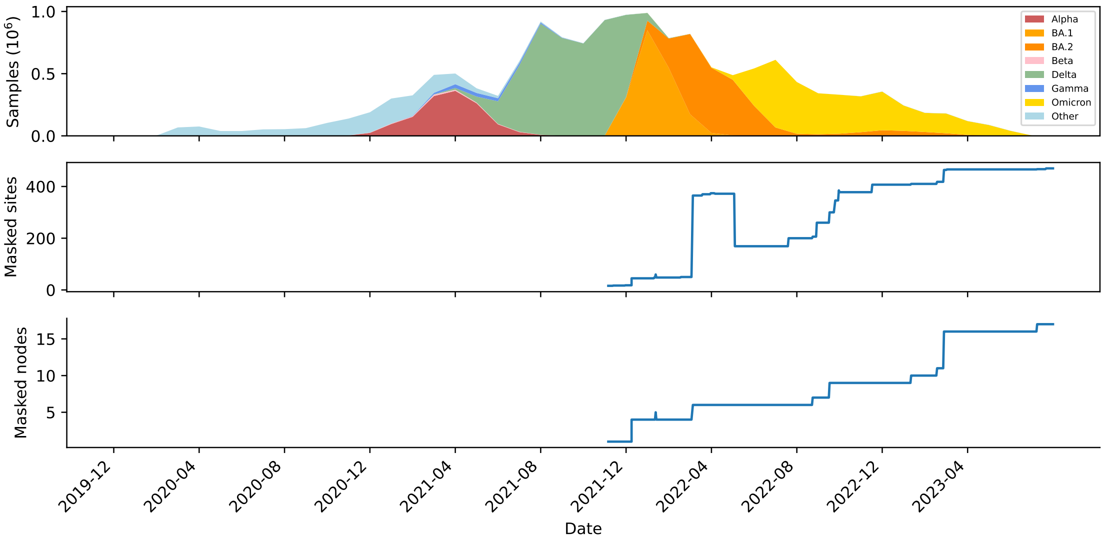


**Figure S1:** Timeline of the SARS-CoV-2 pandemic from December 2019 to July 2023, plus the number of masked sites and nodes in the SARS-CoV-2 global phylogenetic tree.

#### Primer scheme identification validation

Example scheme score plots output by Viridian for a very clean Illumina sample (ERR9362110) and an Illumina sample with fragmented reads (ERR8959211) are shown in Supplementary Figure S2. These are from the truth dataset, and are typical of those samples: the ARTIC version 3 Ilumina reads (eg ERR8959211) are fragmented due to tagmentation during library preparation, whereas the ARTIC version 4 reads are not. Artemis screenshots of the reads from these runs are given in Supplementary Figure S3, showing the difference between the two runs.

ERR9362110 was sequenced using ARTIC scheme version 3, which Viridian scored at 4920. The other scores ranged from -278 to 632. ERR8959211 was sequenced using ARTIC scheme version 4.1, which Viridian scored at 2372. The other scores ranged from -126 to 504. The comparatively lower score of 2372 is a result of the fragmented reads, but is still 4.7 times greater than the second best score. This shows that Viridian successfully calls the scheme even when the reads within each amplicon are fragmented.

The accessions of the manually checked runs that were discordant between the ARTIC primer scheme version in the INSDC metadata and the Viridian call were:

- Illumina, INSDC=3, Viridian=4: ERR7207071, ERR7687763, ERR7696315, ERR7704807, ERR7713199
- Illumina, INSDC=4, Viridian=3: ERR6435020, ERR7202077, ERR7306912, ERR8190486, ERR8228569
- Nanopore, INSDC=3, Viridian=4: ERR5226357, ERR8202943, ERR8218048, ERR8235241, ERR8250042
- Nanopore, INSDC=4, Viridian=3: ERR5226357, ERR5401980, ERR5516251, ERR6114066, ERR6207127.

All Nanopore runs followed the same pattern: reads mapped at positions corresponding exactly to complete amplicons, and all matched the scheme version called by Viridian. Artemis screenshots of Nanopore run ERR5226357 are shown in Supplementary Figure S4. The Illumina reads were fragmented, but with enough signal to determine that the Viridian call was correct in 9 of the 10 runs, and the remaining run ERR8228569 was inconclusive. Artemis screenshots of Illumina run ERR7704807 are shown in Supplementary Figure S5, which is typical of the 9 runs whose scheme was manually identified. The inconclusive run ERR8228569 is shown in figure S6.

The screenshots focus on the last $\sim$10kb of the genome, since this is where the amplicons differ most between the two scheme versions. Reads were randomly sampled using SAMtools with the `-s` option before viewing to aid visualisation, since the full depth results in stacks of reads being too high for the viewing window and therefore not visible.

(a)

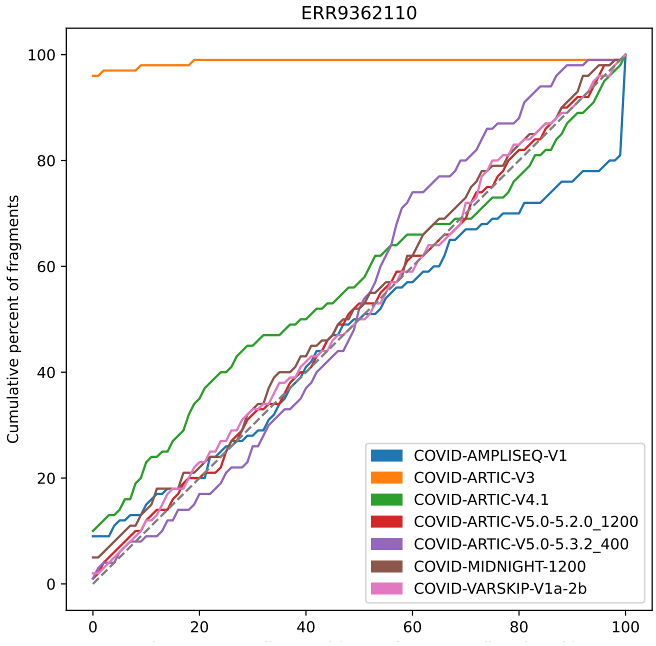


(b)

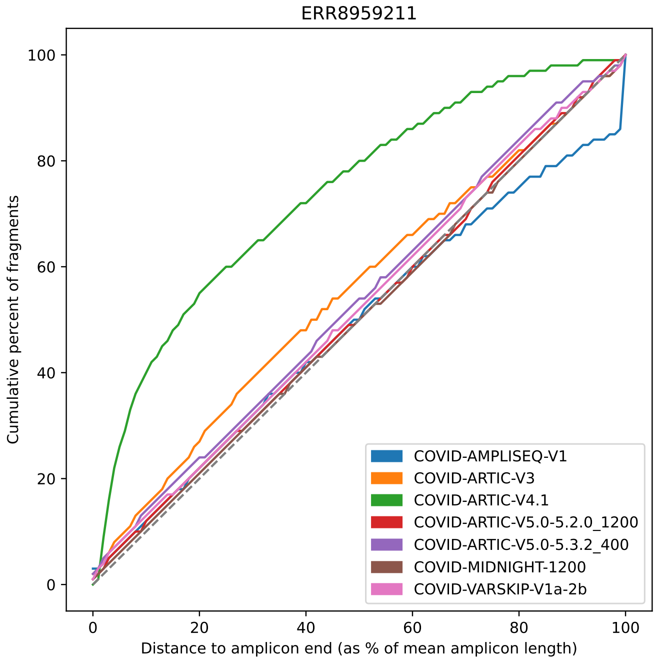


**Figure S2:** Scheme identification plot output by Viridian for Illumina runs (a) ERR9362110 and (b) ERR8959211.

(a)

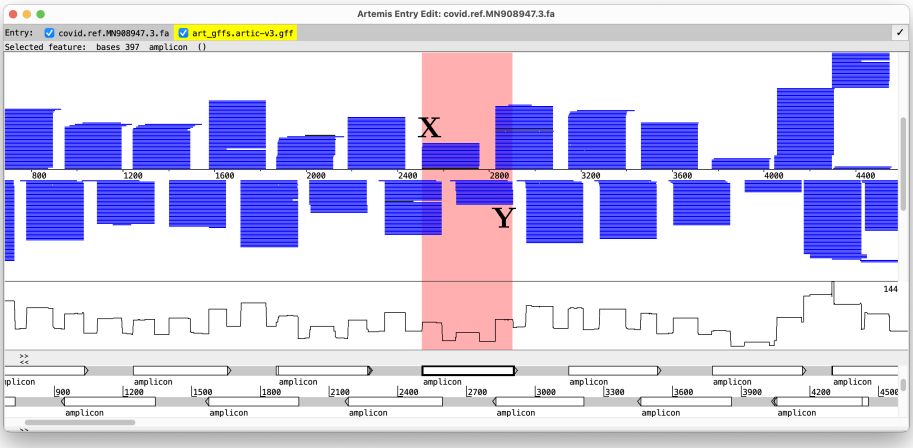


(b)

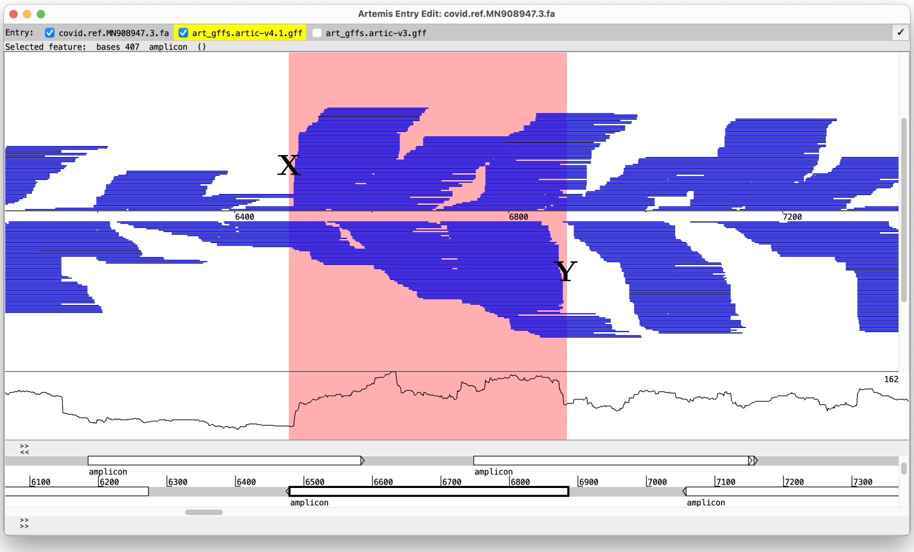


**Figure S3:** Artemis screenshots showing reads mapped to the SARS-CoV-2 reference genome: (a) ERR9362110; (b) ERR8959211. Reads are shown using the "strand stack" view, where the upper reads are those that map in the forwards orientation, and the lower reads are those mapped in the reverse direction (flag 16 in the BAM file). The line plot below the reads shows the read depth across the genome. Since amplicons overlap, they are shown as annotated alternating between the forward and reverse strands. This is to aid visualization and the apparent strand/direction of each amplicon is not relevant. It is their positions that is important. An amplicon shown on top of another amplicon is where there are alternative primers for the same amplicon, for example (a) at position ∼4,400. One amplicon is highlighted in each screenshot to illustrate how the ends of mapped reads match. We are looking for reads mapped to the forwards strand with left ends matching the amplicon start (marked with an X), and reads mapped to the reverse strand with right ends matching the amplicon end (marked with a Y): in (a) they match perfectly, in (b) there is enough of a signal to see that the reads match, but is less clear.

(a) (b)

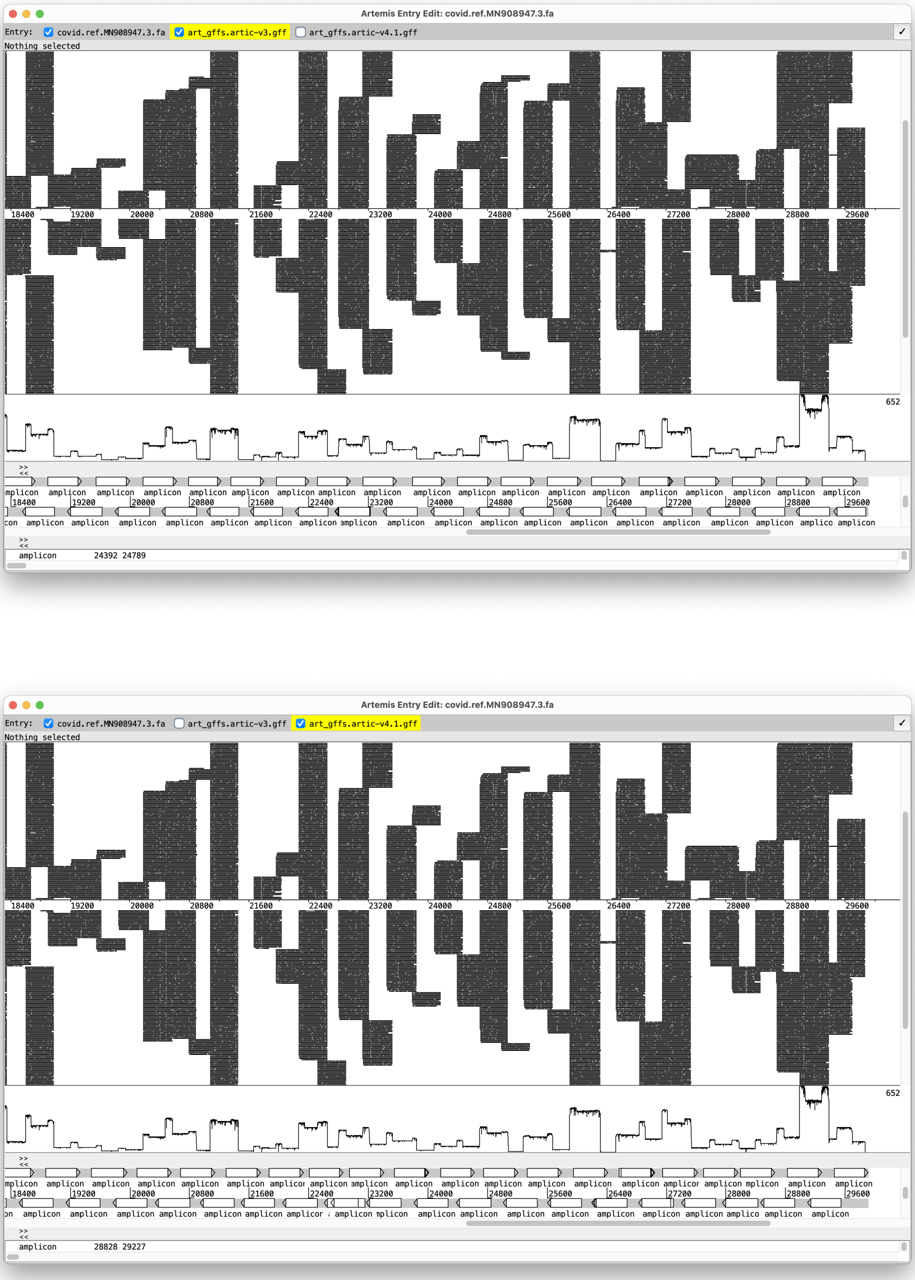


**Figure S4:** Artemis screenshots showing reads from Nanopore run ERR5226357 mapped to the SARS-CoV-2 reference genome. The screenshots are identical, except for the lower track showing the amplicons from ARTIC primer scheme version 3 in (a) and version 4 in (b). Reads are shown using the “strand stack” view, where the upper reads are those that map in the forwards orientation, and the lower reads are those mapped in the reverse direction (flag 16 in the BAM file). The line plot below the reads shows the read depth across the genome. Since amplicons overlap, they are shown as annotated alternating between the forward and reverse strands. This is to aid visualization and the apparent strand/direction of each amplicon is not relevant. It is their positions that is important. An amplicon shown on top of another amplicon is where there are alternative primers for the same amplicon, for example (b) at position ∼22,700. The reads match perfectly to scheme version 3.

(a)

(b)

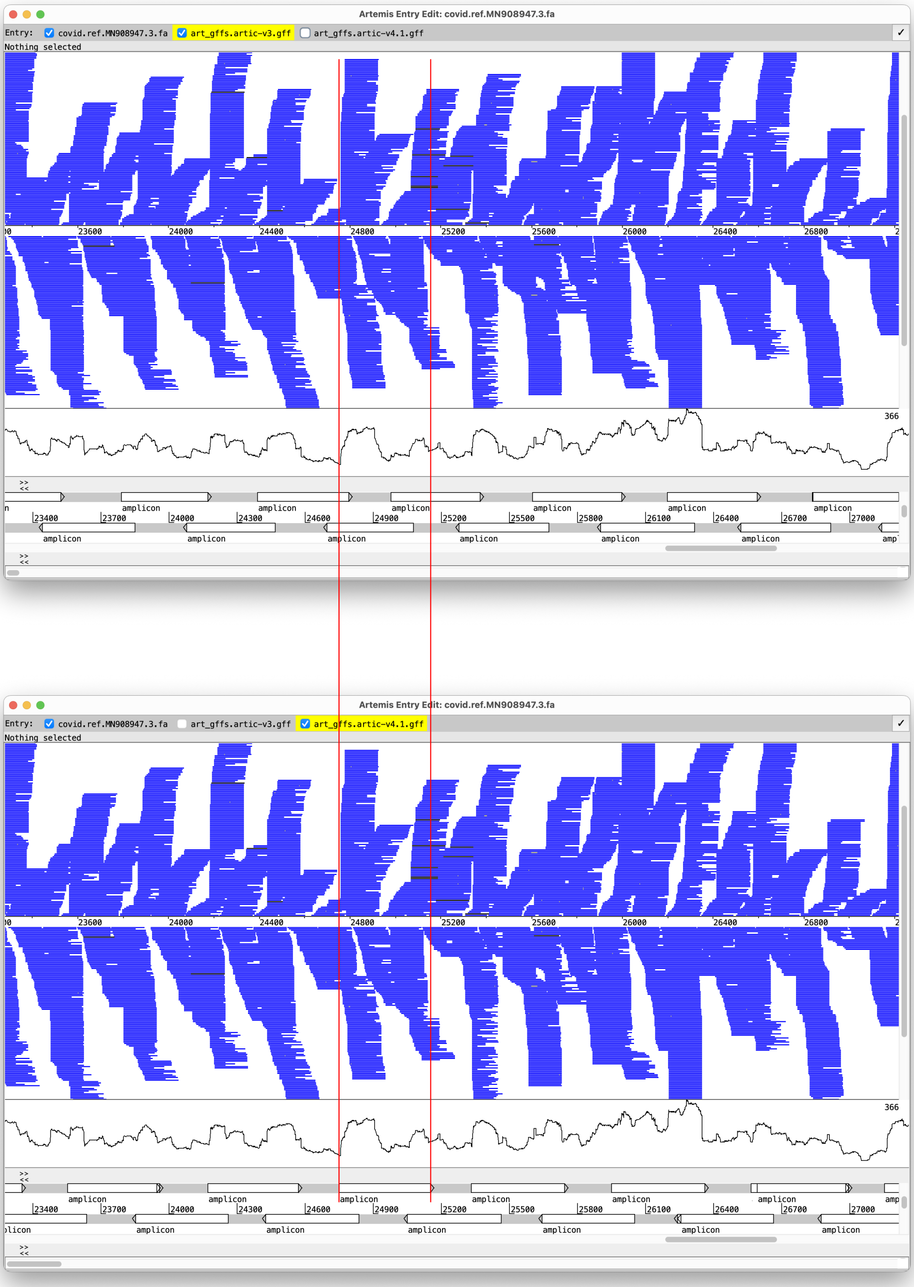


**Figure S5:** Artemis screenshots showing reads from Illumina run ERR7704807 mapped to the SARS-CoV-2 reference genome. (a) ARTIC amplicon scheme version 3 is annotated. (b) ARTIC amplicon scheme version 4 is annotated. See the legend of Supplementary Figure S5 for an explanation of the visualisation details. The reads best match scheme version 4: large increases/decreases in read depth match the start/end of amplicons, and there are peaks of greater read depth where adjacent amplicons overlap. See for example the amplicon marked by the two vertical red lines.

(a)

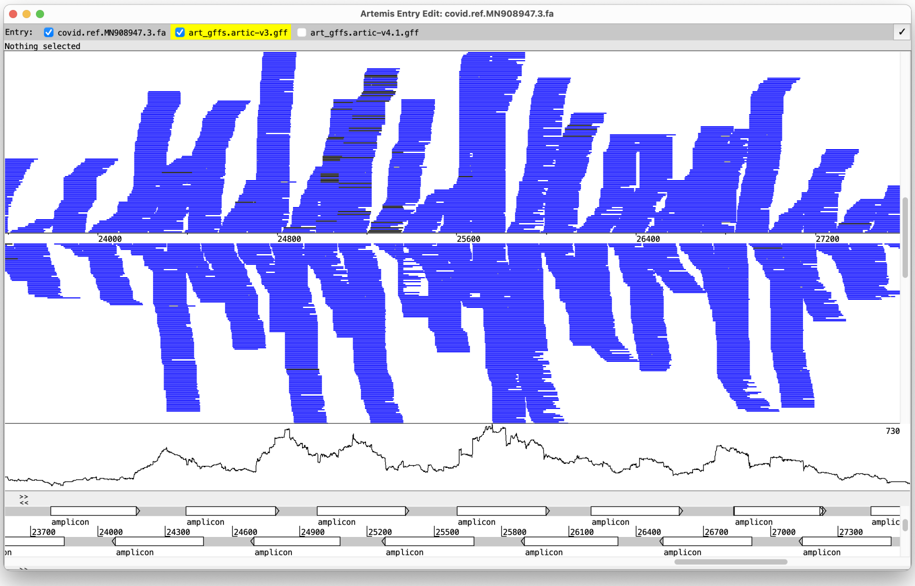


(b)

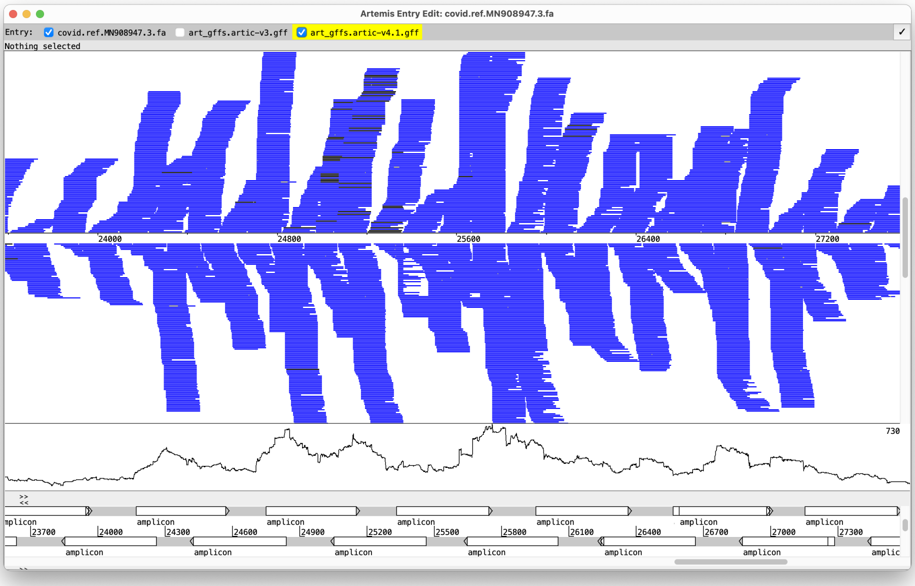


**Figure S6:** Artemis screenshots showing reads from Illumina run ERR8228569 mapped to the SARS-CoV-2 reference genome. (a) ARTIC amplicon scheme version 3 is annotated. (b) ARTIC amplicon scheme version 4 is annotated. See the legend of Supplementary Figure S6 for an explanation of the visualisation details. For this Illumina run, there is no clear match to either amplicon scheme.

#### Run time and memory

A summary of the run time and memory usage on the truth dataset is shown in figure S7. Values are taken from the output of the Unix command `/usr/bin/time`. Plots generated from the full results in Supplementary Table S6.

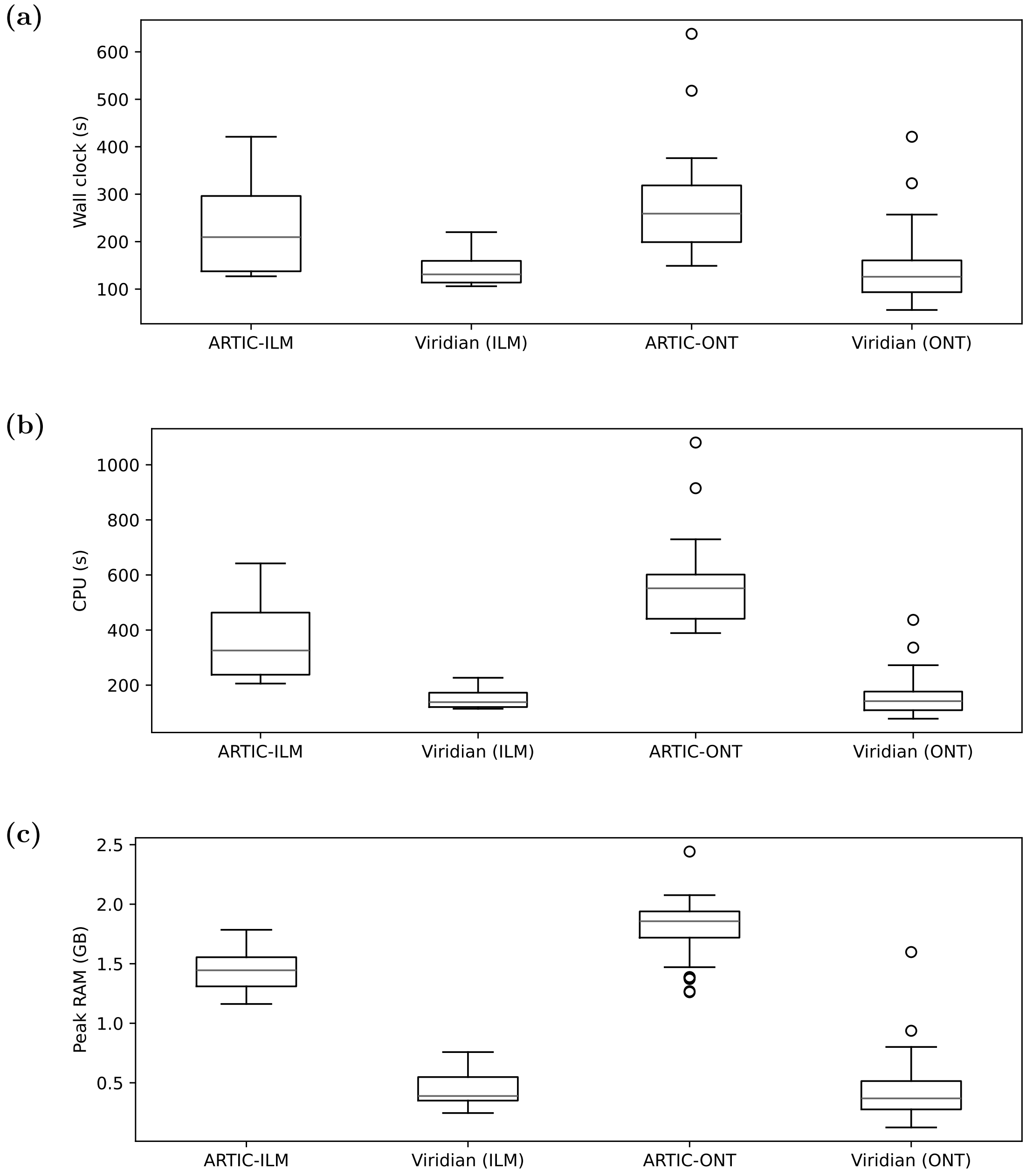


**Figure S7:** Comparison of a) wall clock time, b) total CPU time, and c) peak RAM usage on the truth dataset. Viridian results are split into Illumina and ONT, for comparison with the separate pipelines ARTIC-ILM and ARTIC-ONT.

#### Indel calls

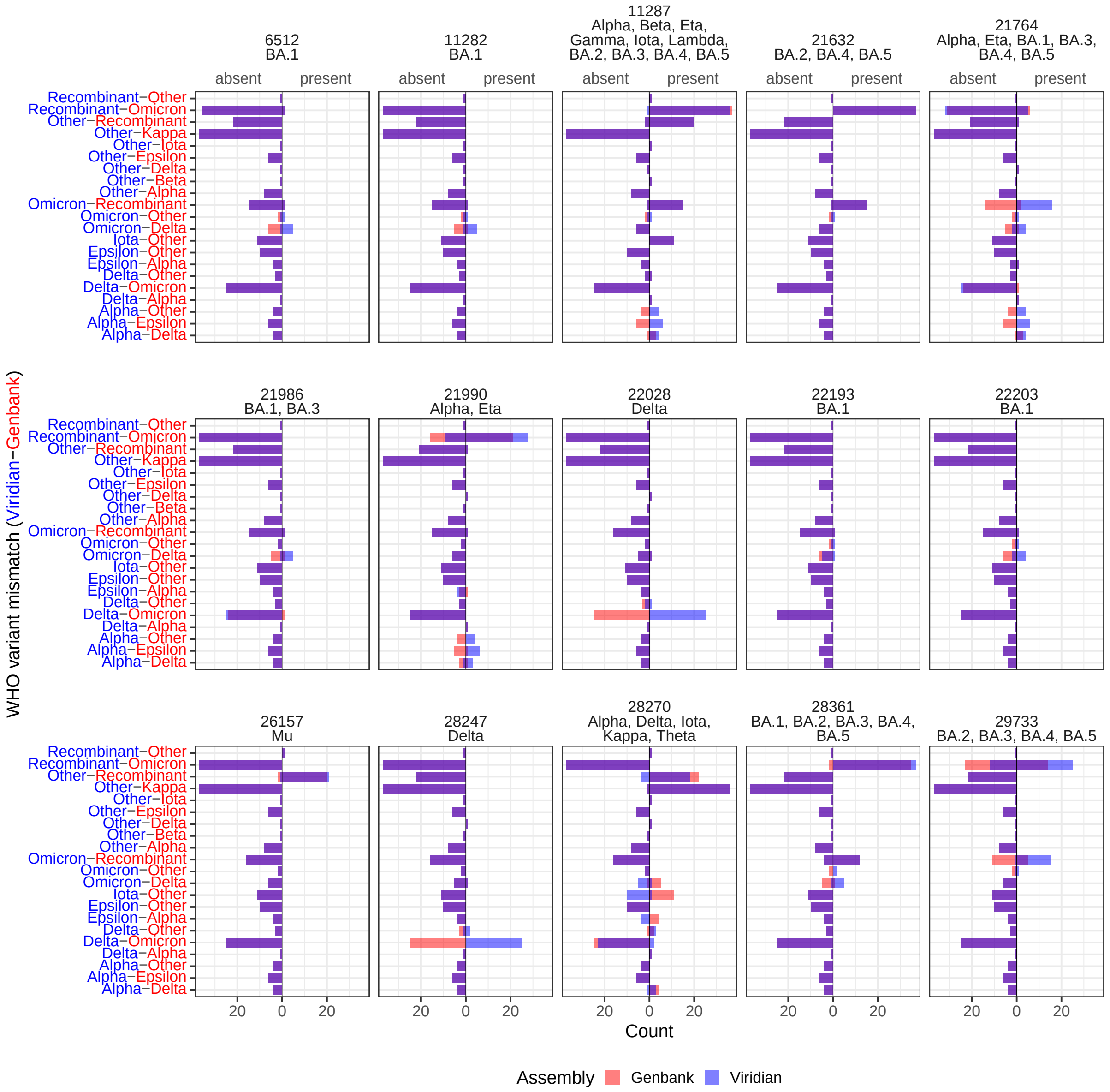


**Figure S8: VOC-defining indels in samples where Viridian and GenBank disagree on Pango assignment.** For (the few) genomes where the Pango WHO variant-of-concern assignment differed between Viridian and GenBank, for each defining indel within an official variant consensus, we compared the number of samples where the indel was not identified (left of black line) to that where it was (right of black line) using Viridian (blue) and Genbank (red). The purple bar overlap shows where the presence/absence is consistent between the two assemblies. The WHO variants in which the indel is consensus are listed under the site identifier. Overall the results are very consistent, with the biggest discrepancies being where Viridian identifies Delta-defining indels and the sample is called as Delta, whereas GenBank does not not call the indel, identifying the sample as Omicron.

### Reversions

**a** **b**

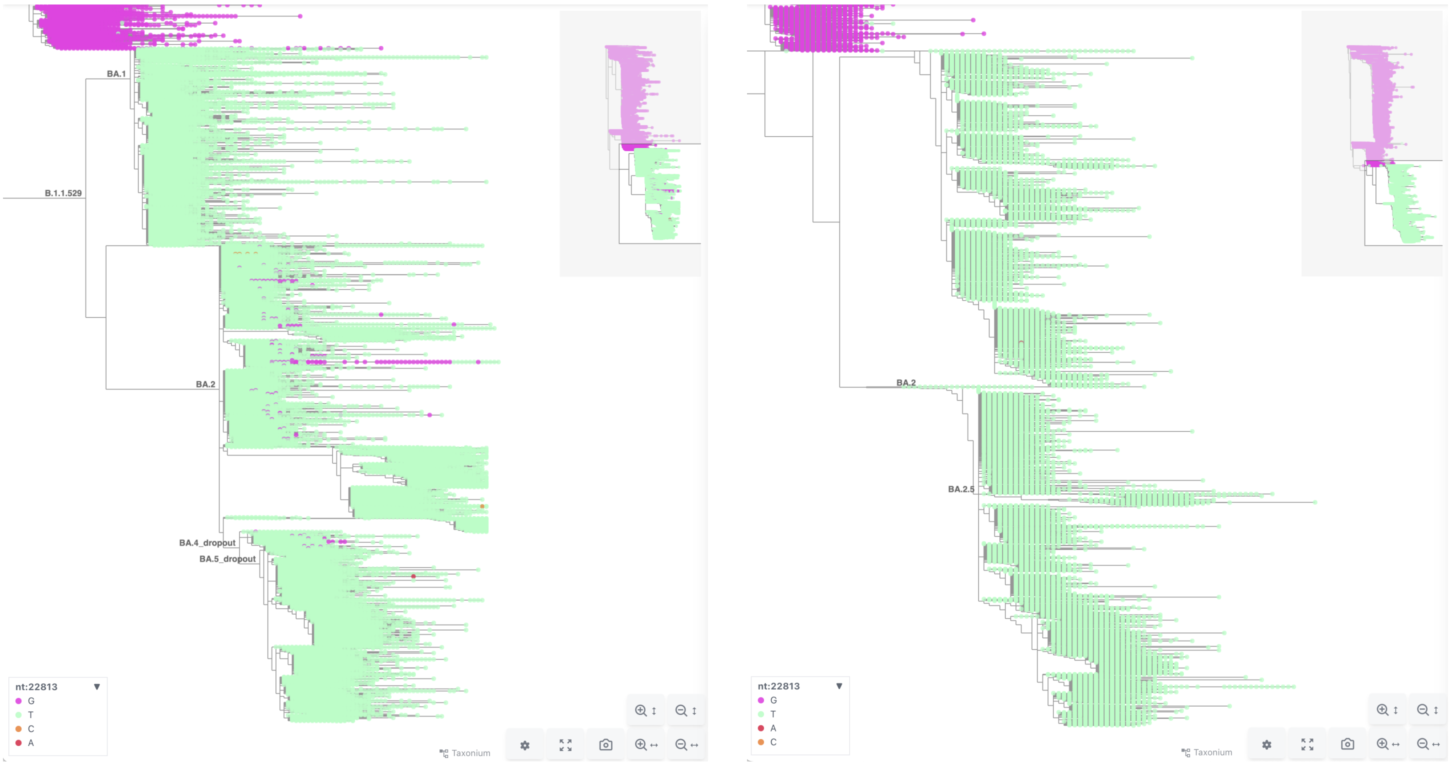


**Figure S9:** Taxonium screenshots of SARS-CoV-2 phylogenies, coloured by genotype at genome position 22813 (spike codon 417). a) The current UShER global phylogeny. b) The global Viridian phylogeny. Samples with the ancestral/reference genome allele are pink, and other genotypes (nearly all green) are shown in other colors.

#### Improved accuracy of lineage growth rate estimate

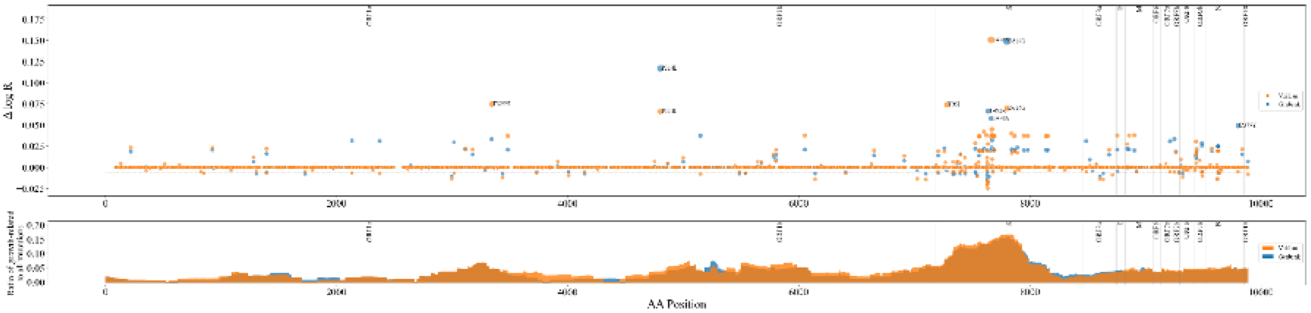


**Figure S10:** Manhattan plot showing mutation relative growth rate $\Delta \log R$ ($y$ axis) by position ($x$ axis) of mutations across the genome for each dataset, with reading frame annotated above. Relative growth rate $\Delta \log R$ is the contribution by a given mutation to the common log of the growth rate of a mutated strain divided by the growth rate of the ancestral strain. The 5 highest-growth mutations from each dataset are annotated. The standard deviation of mutation growth rates across both datasets is 0.006 – dotted lines at $\pm 0.006$ are drawn to indicate growth-related mutations (mutations with $|\Delta \log R| > 0.006$). (b) The ratio of count of growth-related mutations to count of all mutations within a 600-amino-acid width window of $x$ axis position is shown. Fisher's Exact Test is performed on the count of growth-related and non-growth-related mutations in each reading frame, with no statistically significant differences observed.

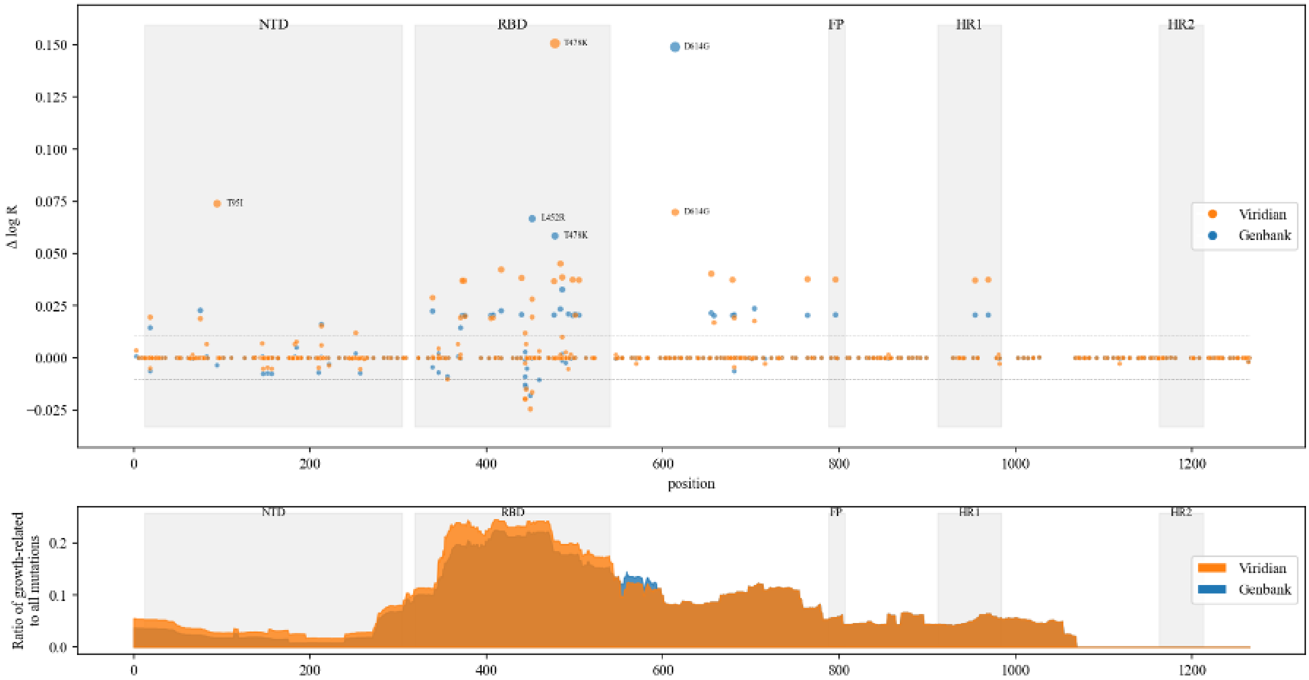


**Figure S11:** Mutation relative growth rate $\Delta \log R$ ($y$ axis) by position ($x$ axis) of mutations in the spike protein for each dataset, with 3 highest-growth mutations from each dataset annotated. Notably, when switching from GenBank to Viridian data, the growth rate of D614G approximately halves while the growth rate of T478K approximately doubles. (b) Ratio of count of growth-related mutations to count of all mutations within a 200-amino-acid width window of $x$ axis position is shown. Each subregion (N-Terminal Domain (NTD), Receptor Binding Domain (RBD), Fusion Peptide (FP), Heptad Repeats 1 and 2 (HR1 and HR2)) is shaded and Fisher's Exact Test is performed for difference in proportions, yielding no statistically significant differences.

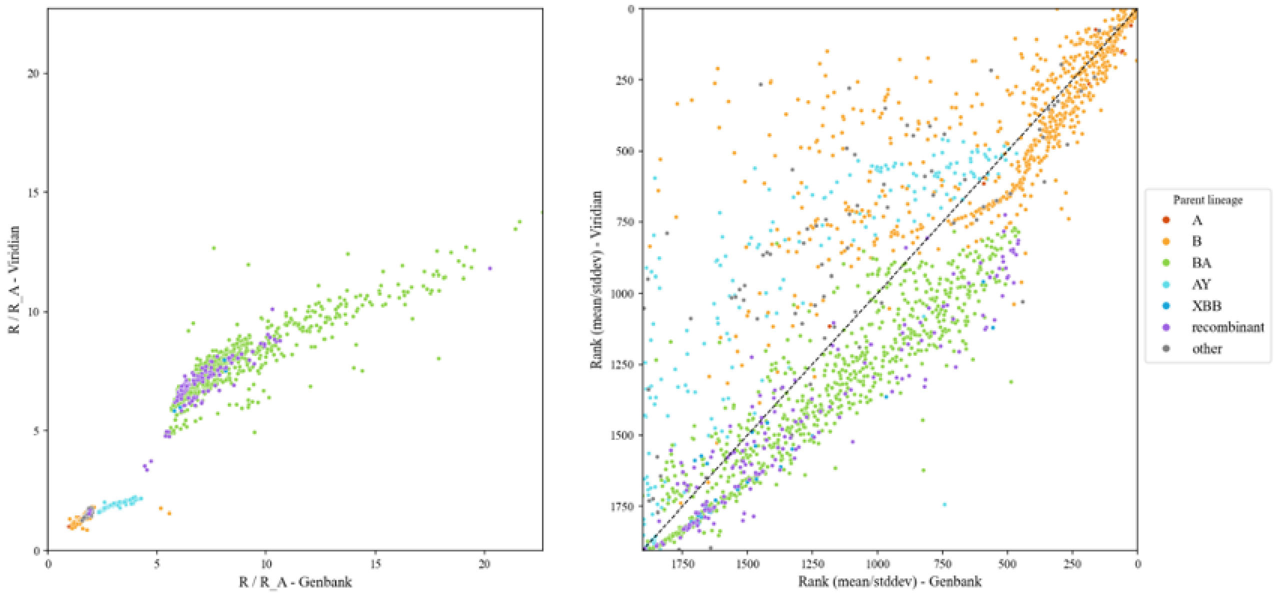


**Figure S12:** Note: legend labels denote parent lineage. (a) Relative growth rate of strain using Viridian data ($y$ axis) versus GenBank data ($x$ axis). Both datasets yield the result of growth rate clustering into two major clouds, mostly categorized by emergence of BA and recombinant lineages (and their sub-lineages). While we don't expect relative growth rate $R/R_A$ to be exactly preserved across datasets (due to a different number of mutations, etc.), we do expect relative order to be consistent. (b) Rank of strain using Viridian data and GenBank data, where rank is determined by mean divided by standard deviation of growth rate posterior distribution. The dotted line $y = x$ is shown. Due to lower uncertainty estimates a posteriori using the Viridian data, there is a frequent shift of strains with poor rank using the GenBank data having better rank using the Viridian data, especially among B lineages and AY sub-lineages. This mean/stddev metric is common for feature selection, among other tasks. Since figure 4(a) shows that there is not much change in rank of mean $R/R_A$, we can attribute most of the changes in rank (mean/stddev) to changes in stddev. The points that lie above the y=x line are those for which uncertainty in the standard deviation of the $R/R_A$ estimate likely decreased. This shows the power of Viridian in helping to decrease uncertainty values and prioritize different strains (notably AY and B) compared to GenBank.

#### Impact on evolutionary and epidemiological analysis

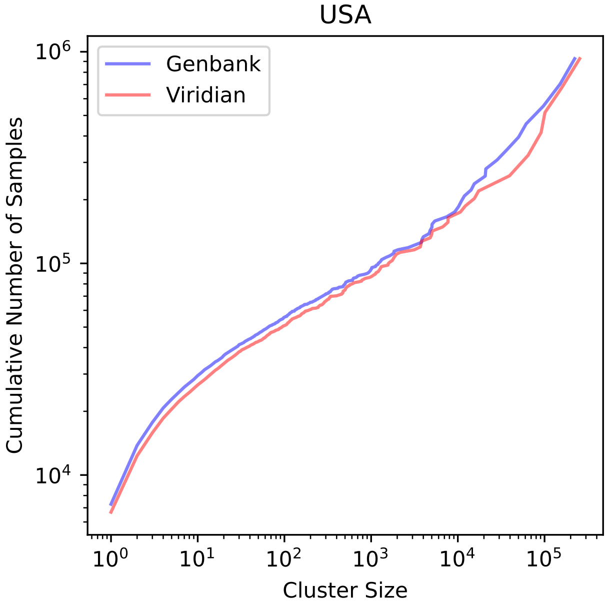


**Figure S13:** Cumulative Distribution of the number of samples in USA stratified by cluster size.

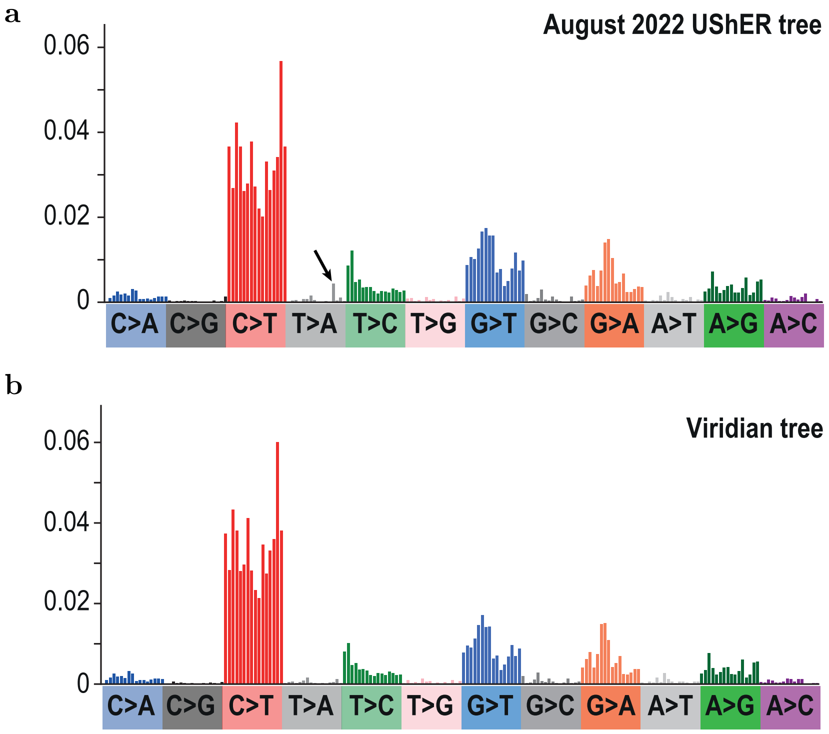


**Figure S14:** Comparison of Alpha variant mutational spectra calculated using (a) the August 2022 UShER tree [**?**] and (b) the Viridian tree. Colours show different mutation types (for example C mutating to T, labelled as C>T) and bars show individual surrounding contexts (for example an upstream A and a downstream A). Spectra are rescaled by the availability of the starting nucleotide triplet. The arrow shows a contextual mutation that is unexpectedly elevated in the August 2022 UShER tree; this elevation is not present in the Viridian tree.

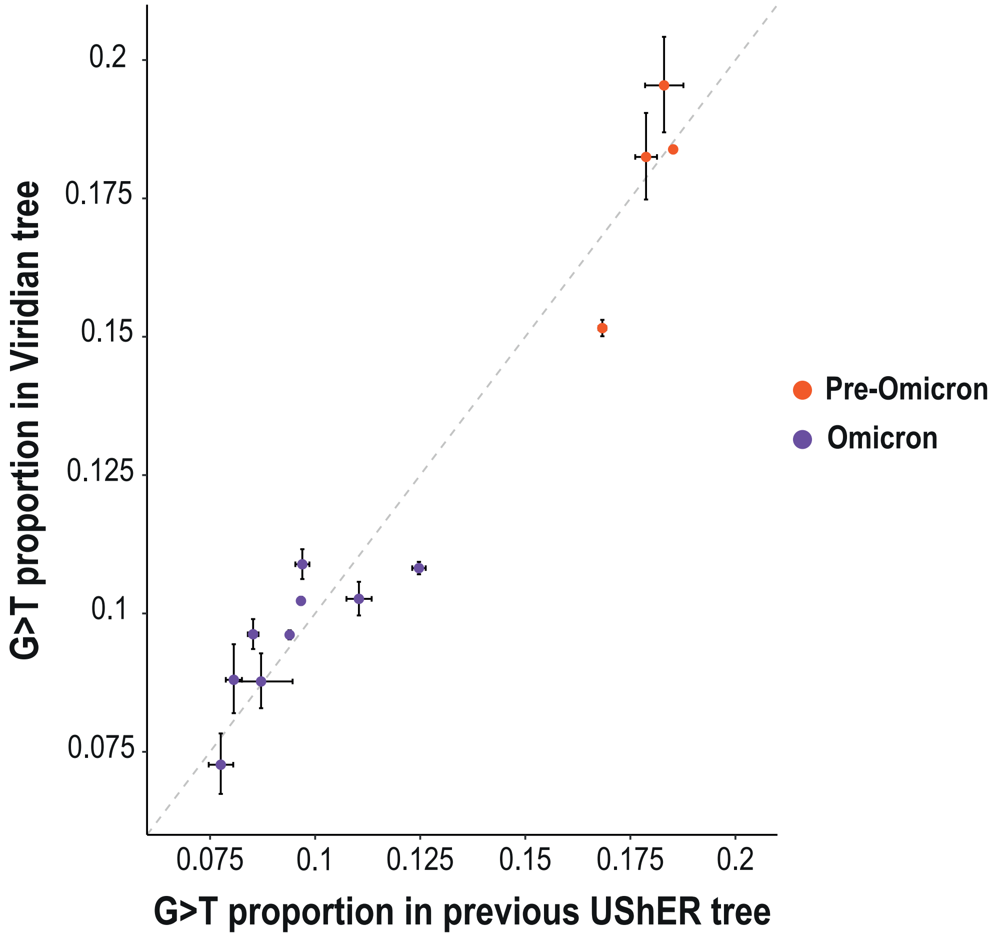


**Figure S15:** Comparison of the proportion of G>T mutations in Omicron and pre-Omicron SARS-CoV-2 lineages between previous UShER trees and the Viridian tree. Points show the proportion of G>T mutations and error bars show the Wilson score interval considering the calculated G>T proportion and number of sampled mutations. A previously observed reduction in G>T mutations in Omicron lineages [Ruis 2023] is still present in the Viridian tree. The date of the previous UShER tree depends on the lineage: August 2022 for Alpha, Beta, Gamma, Delta, BA.1, BA.2, BA.4 and BA.5; October 2023 for BA.2.12.1, BA.2.75, BQ.1, CH.1.1 and XBB.1.5.

#### Geographical distribution of samples

The country for each sample was determined from the "Country" entry in the ENA metadata. The global Viridian tree produced in this study included all INSDC data up to 19th March 2024. The counts of samples for all countries are in Supplementary Table S8, the worldwide and Europe counts are shown in Supplementary figures S16 and S17.

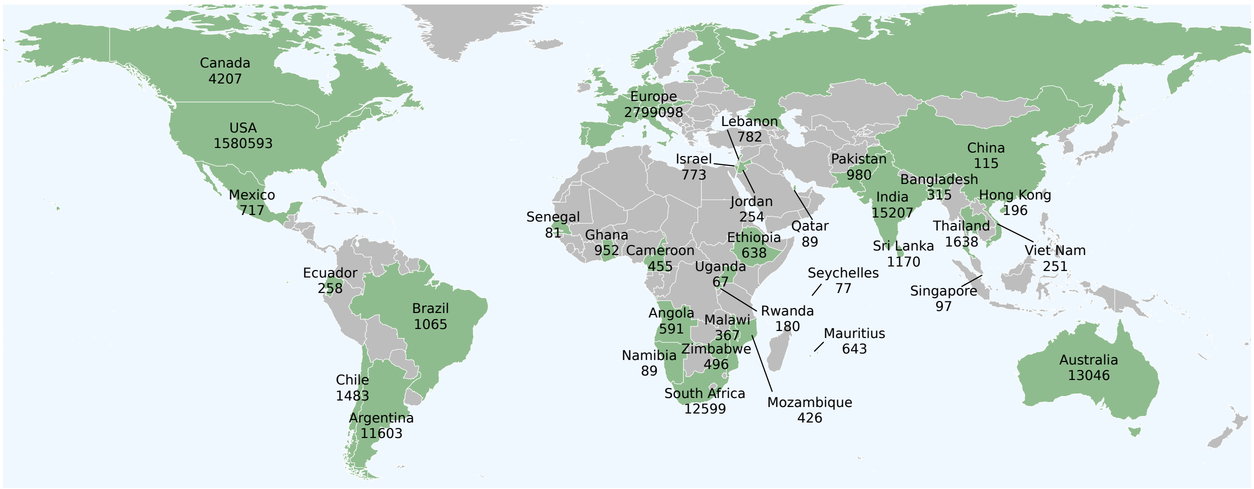


**Figure S16:** Worldwide geographical distribution of samples. Numbers show the total number of samples for each country, excluding QC failures, that are in the global Viridian tree. Only countries with at least 50 samples are labelled, and are coloured in green. See Supplementary Figure S17 for the per-country counts of Europe.

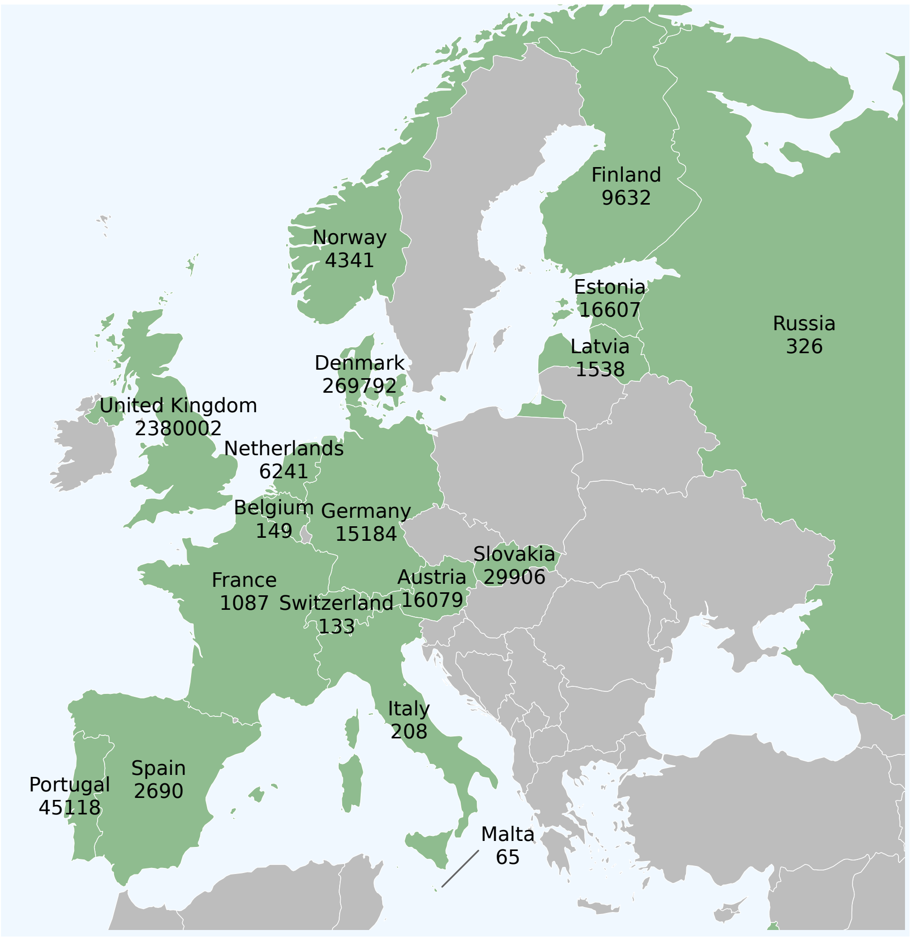


**Figure S17:** Geographical distribution of European samples. Numbers show the total number of samples for each country, excluding QC failures, that are in the global Viridian tree. Only countries with at least 50 samples are labelled, and are coloured in green. See Supplementary Figure S16 for worldwide counts.
